# Promoter hijacking by primate LINC00473 disrupts an ancestral CREB-PDE10A feedback loop

**DOI:** 10.1101/2025.10.02.680088

**Authors:** Shelby J. Squires, Brandon S. Smith, Mukhayyo Sultonova, Emma J. Campbell, V. Shruthi Bandi, Brandon I. MacDonald, Joao A. Paulo, Adam P. Johnston, J. Patrick Murphy, P. Joel Ross

## Abstract

Thousands of human long noncoding RNAs (lncRNAs) evolved in primates, although much remains unknown about how these lncRNAs shaped human gene regulatory networks. The primate-specific lncRNA gene *LINC00473* has a CREB-inducible promoter that is conserved beyond primates and the gene is located upstream of PDE10A, which encodes an inhibitor of CREB. To gain insight into the regulatory consequences of *LINC00473* acquisition, we tested the cellular function of the conserved mouse promoter. We found that the homologous mouse promoter is induced by CREB and regulates the downstream *Pde10a* gene in C2C12 myoblast-like cells and neurons. Activation of this promoter by CRISPRa increased *Pde10a* transcript and protein levels, and proteomics revealed that elevated Pde10a promoted C2C12 differentiation at the expense of proliferation. Activation of *Pde10a* by CRISPRa also impaired CREB-dependent gene expression, suggesting that the mouse homolog of the *LINC00473* promoter drives a CREB-Pde10a feedback loop. In contrast to the mouse homolog, human *LINC00473* promoter activation by CRISPRa increased *LINC00473* expression with either no change in *PDE10A* or delayed induction compared to mice. Our findings suggest that the newly evolved *LINC00473* gene hijacked an ancestral CREB-inducible *Pde10a* promoter, thereby disrupting a negative feedback loop that may otherwise constrain CREB-dependent gene expression.

**GRAPHICAL ABSTRACT:** 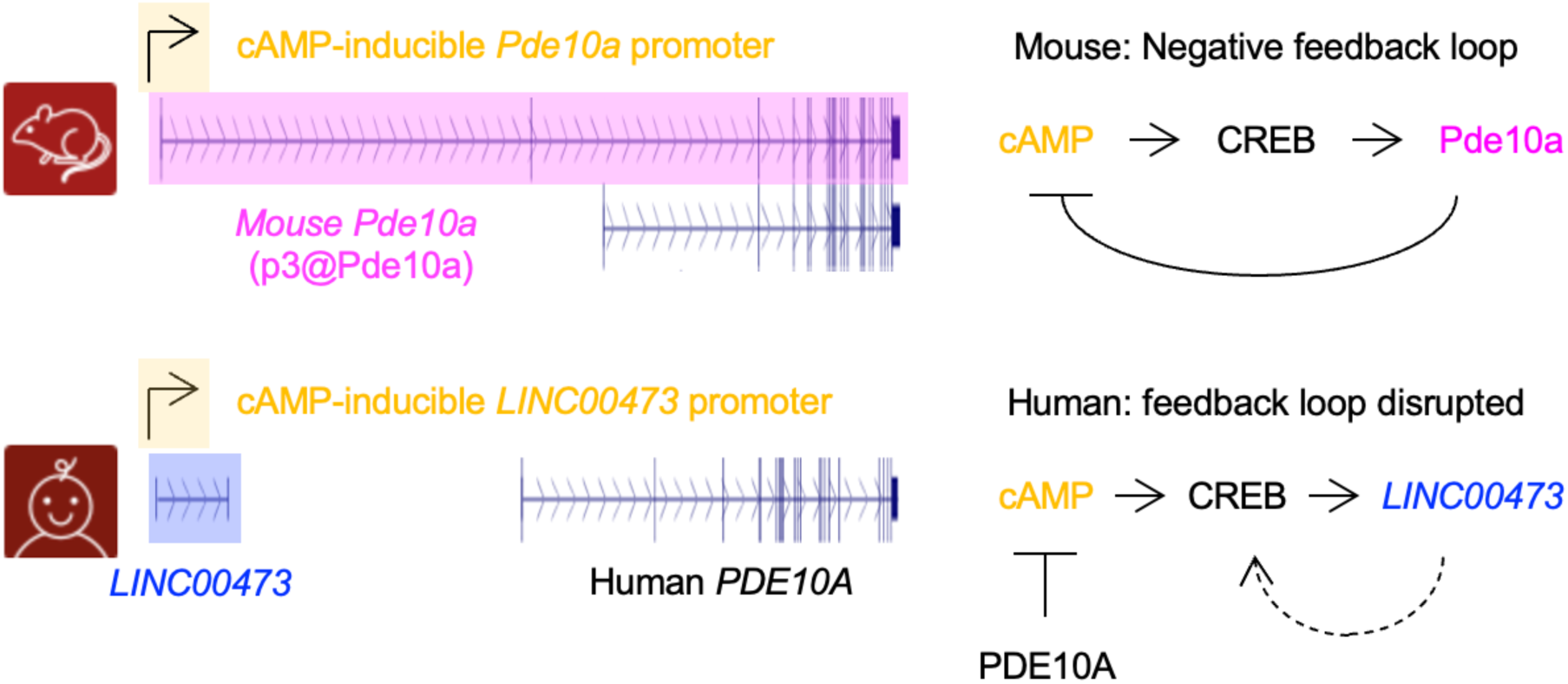

## INTRODUCTION

Despite striking differences in size, morphology, and cognition, humans share ∼99% sequence identity with our nearest relative the chimpanzee (1) and 85% sequence identity with the widely used mammalian model system the laboratory mouse (2). The most striking genomic differences between humans and other mammals are in the noncoding genome: vast swaths of repetitive “junk” DNA (3) punctuated with regulatory elements and noncoding RNAs, which together regulate how protein-coding exons of the genome are expressed (4–7). Of the regulatory elements of the noncoding genome, long noncoding RNAs are a particularly promising avenue for exploring the evolution of human complexity. Humans encode approximately 11, 000 primate specific lncRNAs (8), compared to only ∼80 primate-specific protein coding genes (9), and much remains unknown about the roles of newly evolved lncRNAs in human gene regulatory networks.

lncRNA genes cooperate with DNA regulatory elements like promoters and enhancers to direct and fine-tune the expression of protein coding genes (4–7). lncRNAs are defined as transcripts of at least 200 nucleotides that have little or no capacity to encode functional polypeptides (4). The emerging consensus about lncRNA function is that they regulate expression of other genes, including protein coding genes, via RNA-dependent or RNA-independent mechanisms (5, 7, 10). The RNA products of lncRNA genes can interact with specific gene regulatory proteins and recruit them to target genes or transcripts. lncRNAs may also influence expression locally via the lncRNA’s cis regulatory elements or via transcription-associated processes. lncRNAs evolve much more rapidly than protein coding genes, resulting in lncRNAs being relatively restricted to specific species or evolutionary lineages compared to protein coding genes. Notably, lncRNA promoters are typically much better conserved than their exons, and exhibit similar conservation levels compared to promoters of protein coding genes (11). This has led to the widely held notion that for many lncRNAs it is the act of transcription itself, rather than the sequence of the resultant lncRNA, that is of primary functional importance (4). However, another possibility is that some lncRNAs co-opted existing promoters or enhancers (12) that had distinct functions prior to evolutionary acquisition of the lncRNA exons.

*LINC00473* is a lncRNA that first evolved in a common ancestor of monkeys, apes, and humans, and is thought to play roles in neuropsychiatric disorders and higher order brain functions (13–15). Like some other annotated lncRNAs, human *LINC00473* encodes a polypeptide (16, 17), although the open reading frame has been lost due to mutation in other closely related primates like chimpanzees and it is thought to play a negligible role in *LINC00473* function (13). *LINC00473* expression is dynamically regulated by conserved signaling pathways that activate CREB – a central regulator of neuroplasticity (18) – including synaptic activity and cyclic AMP (cAMP) (13, 16, 19, 20). Ectopic expression of human *LINC00473* in mouse neurons, which otherwise lack this lncRNA, enhances CREB-dependent transcription and increases neuronal excitability (13). Although mechanisms remain unclear, *LINC00473* may promote interactions between the CREB coactivators NONO and CRTC1 (21) and nuclear retention of CRTC1 (13). In the context of neuropsychiatric disorders, *LINC00473* was shown to be downregulated in the medial prefrontal cortex of females with depression, but not depressed males (14). Consistent with this, transgenic gain-of-function studies in mice revealed sex-dependent behavioral phenotypes, with viral vector-mediated ectopic expression resulting in sex-specific improvements in stress resilience (14) and neuroeconomic decision making (15) in female mice. These findings show that a primate-specific lncRNA can influence CREB-dependent gene expression, neural excitability, and behavioral outcomes to have physiological RNA-dependent functions, even in species that lack *LINC00473*.

Many lncRNAs regulate neighboring genes, although the relationship between *LINC00473* and its nearest chromosomal neighbor *PDE10A* (Fig. 1A) remains unclear. *LINC00473* and *PDE10A* functionally converge on cAMP/CREB signaling (22, 23), *PDE10A* encodes a phosphodiesterase that degrades cAMP (24, 25) and dampens the activity of CREB (26) and may therefore oppose LINC00473 function (13, 21). Like *LINC00473*, *PDE10A* has also been implicated in a mood disorder: bipolar disorder (27). *PDE10A* has also acquired some novel features in the primate genome, including newly evolved exons resulting in a primate specific transcript and protein isoforms (27). *PDE10A* may also have lost an alternative promoter in the primate lineage, coinciding with the acquisition of *LINC00473*. The sequence upstream of *LINC00473* is highly conserved beyond primates, even in rodents, which lack LINC00473 transcripts (13). In the mouse genome, the homology of the *LINC00473* promoter appears to regulate a long *Pde10a* transcript variant (Fig. 1A). Some genome annotations suggest that the human genome encodes similar transcripts arising from the LINC00473 promoter and encoding the PDE10A protein. However, these transcripts predictions (e.g., and XM_017010194.3) result from automated computational predictions that are often incorrect (28, 29). Furthermore, empirical evidence indicates that *LINC00473* and *PDE10A* are independently regulated in human cells (13, 30). Given that *LINC00473* promoter is conserved in species that do not have *LINC00473*, uncovering the ancestral role of this promoter may reveal functional insight into a newly evolved primate lncRNA and uncover unique human gene regulatory processes.

**Figure 1.**
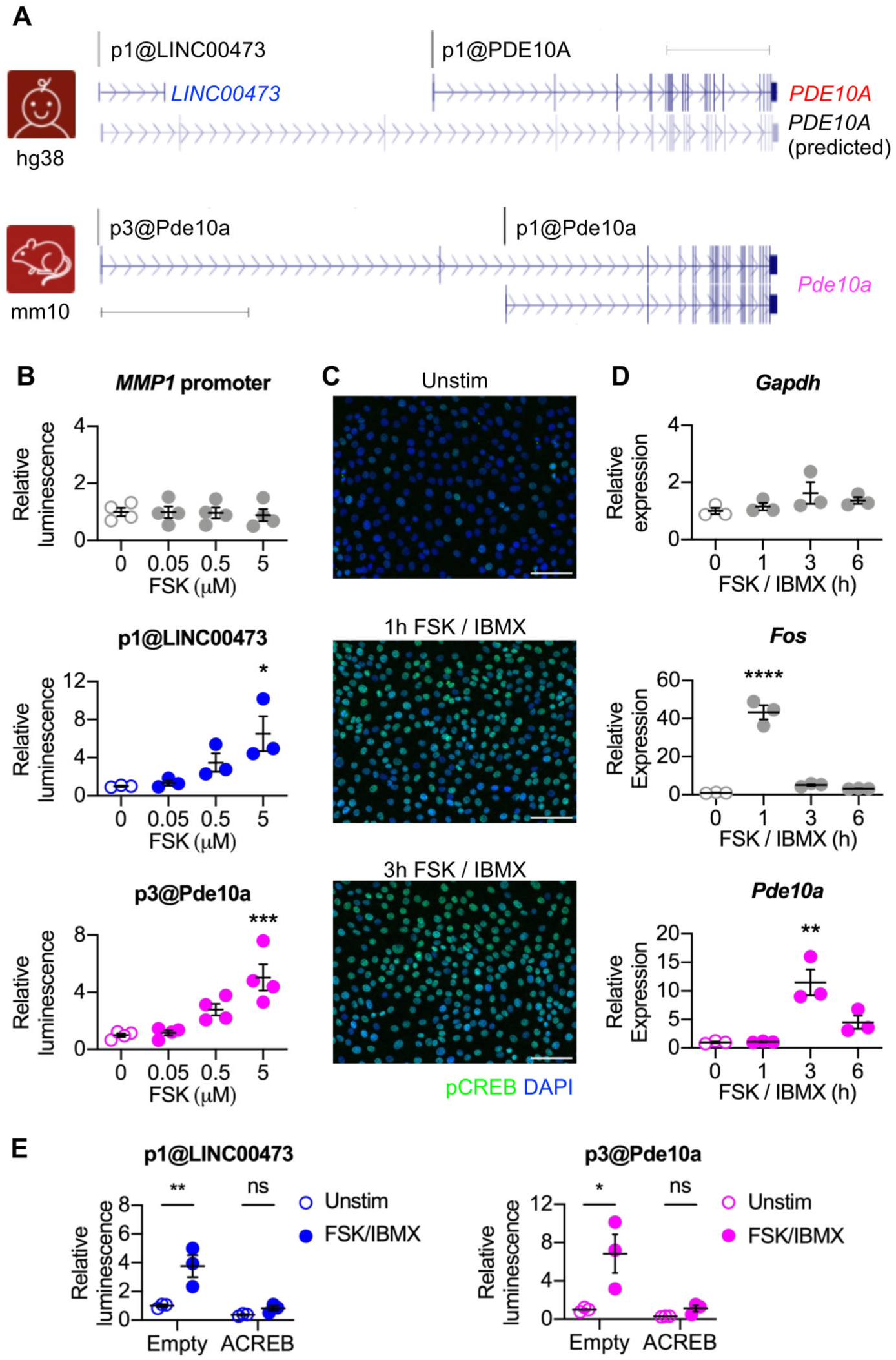
cAMP-inducibility is functionally conserved for the mouse homolog of the *LINC00473* promoter. (**A**) Schematic of human *PDE10A* and mouse *Pde10a* loci. Promoter labels correspond to FANTOM5 CAGE-seq transcriptions start sites (TSS). Schematic of the human *LINC00473* / *PDE10A* genomic locus depicts the location of the CRISPRa gRNA near sgp1@LINC00473, *PDE10A* qPCR primers, and a predicted *PDE10A* transcript startig near sgP1@LINC00473 (XM_011535387.4). Representative transcripts are displayed for each TSS / transcription unit. Modified from UCSC Genome Browser (GRCh38/hg38, GRCm38/mm10); scale bar = 100 kb. **(B)** Reporter assays in 293T cells treated with indicated doses of FSK. Firefly luciferase expression was driven by the *MMP1* promoter (n=4), *LINC00473* promoter (p1@LINC00473; n=3), and mouse homolog of the *LINC00473* promoter (p3@Pde10a; n=4), normalized to renilla luciferase. Data analyzed using one-way ANOVA and Dunnett’s multiple comparisons test. (**C**) Immunocytochemistry to detect CREB phosphorylated at serine 133. Cells were treated for the indicated time with 10 μM FSK and 18 μM IBMX and then fixed for immunocytochemistry. Scale bar = 100 micrometers (**D**) qPCR gene expression analysis in mouse C2C12 cells after 1, 3, or 6 hour stimulation with 10μM FSK and 18μM IBMX normalized to reference genes *Gapdh* / *Rps18* (n=3 biological replicates). Data analyzed using Kruskal-Wallis tests and Dunn’s multiple comparisons test. (**E**) Reporter assays in C2C12 cells transfected with empty vector or vector expressing ACREB (dominant negative CREB) and then left unstimulated or stimulated with 10 μM FSK and 18 μM IBMX Firefly luciferase expression was driven by the *LINC00473* promoter (p1@LINC00473; n=3) or mouse homolog of the *LINC00473* promoter (p3@Pde10a; n=3), normalized to renilla luciferase. Data analyzed using two-way ANOVA and Tukey’s multiple comparisons test. (**B**, **D**, **E**) Dots show individual data points, lines show mean, and error bars show SEM (* p ≤ 0.05; **, P<0.01; ***, P≤0.001; ns, not significant).

Here we investigate the role of the conserved *LINC00473* promoter (Fig. 1A) in regulating cAMP inducible expression of *Pde10a* in the mouse. We demonstrate that cAMP-inducibility is functionally conserved and that activation of the mouse homolog of the *LINC00473* promoter is sufficient to increase *Pde10a* expression. We also explore a potential role for cAMP-inducible *Pde10a* in negative feedback regulation of CREB-dependent gene expression in mouse cells. Our results provide insight into the evolutionary consequence of *LINC00473* acquisition in primates and suggest that *LINC00473* disrupted an ancestral cAMP-driven negative feedback loop.

## MATERIALS AND METHODS

### Ethical approvals

Human induced pluripotent stem cells were previously generated (31) and were cultured for this study under approval of the University of Prince Edward Island (UPEI) Research Ethics Board (REB 6012461). Protein was obtained from mice using procedures that were approved by the UPEI Animal Care Committee in accordance with CCAC guidelines (AUP 16-036).

### Cell culture and pharmacological stimulations

C2C12 (CRL-1772), E14Tg2a (CRL-1821), and MCF-7 (HTB-22) cells were obtained from ATCC via Cedarlane Labs; 293H and 293T cells were obtained from Dr. James Ellis (Hospital for Sick Children, Toronto, ON, Canada). Human induced pluripotent stem cell line PGPC3_75 was kindly provided by Drs. Stephen Scherer and James Ellis (The Hospital for Sick Children, Toronto, ON, Canada). C2C12, MCF-7, 293H, and 293T cells were cultured in high glucose Dulbecco’s modified eagle medium (DMEM) (HyClone SH30022.01) supplemented with 10% fetal bovine serum (FBS) (Gibco 12483020 or Corning 35077CV) and 1% penicillin/streptomycin (HyClone SV30010) and incubated at 37°C with 5% CO2. For passaging, these cells were washed with 1X phosphate buffered saline (PBS) (Corning 21-040-CV) and split using 0.25% trypsin (HyClone SH40042.01). For selection, cells were treated with 1μg/mL (293H) or 2 μg/mL (C2C12) puromycin (Wisent 400-160-EM) and 1mg/mL G418 (Gibco 10131035). For doxycycline (dox) treatment, cells were treated with 0.1-2μg/mL (Sigma D9891-1G). To stimulate cAMP signaling, cells were treated 0.05-10μM FSK (Calbiochem 344270) with or without 18μM IBMX (Sigma I5879-100MG), and MP10 (Sigma PZ0351-5MG) was used at 1-10µM. Culture conditions for mouse embryonic stem cells and human pluripotent stem cells are described below (*Neuronal Differentiation*).

### Gene expression analysis by qRT-PCR

RNA was harvested in 1mL of TRIzol or TRI reagent (ThermoFisher) as described (32), followed by purification using EZ-10 Spin Column Total RNA Miniprep Super Kit (BioBasic BS784) according to the manufacturer’s instructions. For reverse transcription, 300ng total RNA was treated with DNaseI (ThermoFisher EN0521) according to the manufacturer’s instructions. We then proceeded direction to reverse transcription using oligo (dT)18 primers (ThermoFisher SO131) and Maxima H Minus RT (ThermoFisher EP0751), with incubation at 50℃ for 40 minutes, followed by inactivation at 85℃ for 5 minutes.

cDNA was diluted 1:40 to 1:100 for qPCR with SsoAdvanced Universal SYBR Green Supermix (BioRad 1725274) or Luna qPCR mastermix (New England Biolabs M3003E). For each reaction, we mixed 5uL dilute cDNA, 4.5uL SYBR, 0.5uL 2.5uM mix of forward and reverse primers. Primer sequences for qPCR (Table 1) were obtained from PrimerBank and then ordered from ThermoFisher. qPCR was performed using a CFX96 (Bio-Rad) real time PCR system, with thermal cycling conditions consisting of a 3 minute 95°C initial denaturation, followed by 40 cycles of 95°C C for 10 seconds, 60°C for 20 seconds, and 72°C for 20 seconds, followed by a melt curve step. For quantification, the average C_q_ value was determined from three replicate wells and data analysis was performed as described (32). Next, the geometric mean was calculated for the C_q_ value of two reference genes (see figure legends for reference genes). We then calculated ΔC_q_ by subtracting the reference gene mean C_q_ value from the target gene C_q_ value, and log2ΔC_q_ was calculated for each biological replicate. The average log2ΔC_q_ of all biological replicates was calculated for the control group and was used to normalize each value in all experimental conditions. Lastly, we determined the mean of the normalized log2ΔC_q_ for each test group and standard error of the mean.

### Cloning of gRNAs for CRISPRa

To make the gRNA expression plasmids for CRISPRa, we designed and cloned gRNA oligonucleotides into the lentiviral vector plasmid pLCKO (Addgene 73311) as described (33). gRNAs for targeting the human *LINC00473* promoter and the conserved mouse sequence using Benchling software, with all gRNAs having a 5’-NGG-3’ PAM sequence for *S. pyogenes*. For the human sequence, 4 gRNAs were tested and the most effective was used for subsequent experiments (Table 1). The mouse gRNA was chosen because it targets a region in the conserved mouse sequence overlaps a region homologous to the site targeted in by the human gRNA in the *LINC00473* promoter. We also selected a published gRNA targeting the *LacZ* gene to serve as a negative control (34). Cloning of gRNAs done according to published protocols (33) and resultant clones were confirmed by Sanger sequencing at the Centre for Applied Genomics at the Hospital for Sick Children (Toronto, ON, Canada).

### Lentivirus vector infections and CRISPRa Experiments

To generate CRISPRa lentiviral vectors, 293Ts were transfected with lentivirus plasmids and packaging plasmids obtained from Dr. James Ellis (Hospital for Sick Children, Toronto, ON, Canada) (35). *LINC00473*, *Pde10a* or *LacZ* gRNAs were encoded by pLCKO, which also contains a puromycin resistance gene, and dCas9-VP64 was delivered using pHAGE-TRE-dCas9-VP64 (36) (Addgene 50916), which also contains a G418 resistance gene. To package lentivirus vectors, 293T cells in T75 flasks were transfected for 3 hours using branched polyethylenimine (bPEI) in 5% FBS media. For each lentivirus vector, we combined 13.5μg of the lenti plasmid, 4.5μg VSV-G, 9μg Gag/Pol, 9μg REV, with 108μL of 1mg/mL 25kDa bPEI (Sigma 408727) in 1 mL for the pHAGE-TRE-dCas9-VP64 plasmid, we also added 9μg of Tat plasmid (35). Two days after transfection, lenti vectors were collected, filtered with 0.45 micrometer syringe filters, and then aliquoted for storage at −80°C. 293H cells or C2C12 cells were seeded in 6 well plates (∼50, 000 cells per well) and then infected with ∼100-500 μL of each unconcentrated lentiviral in 2mL total of media supplemented with 0.5-8 μg/mL polybrene (Sigma TR-1003). After 3 days, the cells were placed under selection with genetic and puromycin. For CRISPRa qRT-PCR experiments, *LacZ* and *Pde10a* gRNA CRISPRa cells were treated with dox for 6 days, followed by RNA isolation.

### Immunocytochemistry

Cells for fixed using 4% paraformaldehyde (ThermoFisher 43368), permeabilized cells with 0.1% TritonX-100 (BioBasic TB0198) and blocked cells with PBS supplemented with 5% goat serum (Millipore S26-100ML) and 1% bovine serum albumin (ThermoFisher BP1600-100). Fixed cells were incubated overnight at 4℃ in with diluted primary antibody (anti-HA, Invitrogen PI26183, 1:500; anti-phospho-CREB, Santa Cruz Biotechnology sc-81486, 1:200; anti-OCT4, Santa Cruz Biotechnology sc-5279; anti-beta-III-tubulin, Millipore MAB1637, 1:200; anti-Nestin, Aves Labs NES, 1:200; anti-HuC/D, ThermoFisher A21271, 1:250; anti-MAP2, ThermoFisher PA517646, 1:250) in blocking solution. After overnight incubation, cells were washed three times with PBS-T and then incubated cells for 1 hour at room temperature with anti-mouse AlexaFluor-594 or −488 secondary antibodies (Invitrogen A11005, A11008, A78951, A11029, A11012, all 1:500) in PBS-T. After secondary incubation, cells were washed with PBS-T and incubated cells for 5 minutes at room temperature in a 1:5000 dilution of DAPI (Sigma D9542-5MG) in PBS-T.

### Reporter plasmids

Promoters of interest were amplified by PCR using genomic DNA from mouse E14Tg2a or human BJ fibroblasts using OneTaq or Q5 polymerases (New England Biolabs) according to the manufacturer’s instructions. The *LINC00473* promoter (hg38 Chr6:165987785-165988434, reverse strand) includes the FANTOM5 transcription start site peak p1@LINC00473 and the mouse sequence homologous to the *LINC00473* promoter (mm10 Chr17:8524968-8525262, forward strand) includes the FANTOM5 peak p3@Pde10a) (Fig. 1 and Supplementary Fig. S1) (for primers see Table S1). Both the human *LINC00473* promoter and homologous mouse sequences include 2 conserved putative CREB binding sites (Supplementary Fig. S1). These fragments were cloned upstream of firefly luciferase in pmirGLOΔHindIII (a kind gift of Dr. James Ellis) or derivatives with alternative polyadenylation sequences after the firefly luciferase reporter. pmirGLOΔHindIII contains both firefly luciferase (FFLuc) regulated by the promoter of interest and an independently expressed Renilla luciferase (RLuc) gene (driven by the SV40 promoter) downstream of firefly luciferase. To eliminate potential readthrough from the promoter of interest into the RLuc cDNA, we subcloned these constructs into separate plasmids, generating pFFLuc-p1@LINC00473 and pFFLuc-p3@Pde10a, and pRLuc. We also made a derivative of pmirGLOΔHindIII with RLuc regulated by the human phosphoglycerate kinase promoter (pRLuc-PGK). We also generated a control construct pFFLuc construct regulated by the CREB-independent MMP1 promoter (hg38 Chr11:102798099 102798379, negative strand) (Table S1).

### Reporter Assays

For luciferase assays, 293T or C2C12 cells were transfected with reporter plasmids with firefly luciferase driven by inducible promoters of interest and renilla luciferase drive by a control promoter (SV40 or PGK). For initial reporter assay experiments comparing activity between promoters, 293Ts were seeded at 50, 000 cells/well in a 24 well plate. The following day, cells were transfected for 3 hours using 800ng of firefly luciferase plasmid and 200ng renilla luciferase plasmid with 1.5μL Lipofectamine 3000 (ThermoFisher L3000008) and 1μL p3000 reagent following the manufacturer’s instructions. Stimulations started the next day using 0.05, 0.5 and 5μM FSK and continued for 16 hours until cell lysate was harvested in 1X passive lysis buffer (Promega E1941) for reporter assays using the DualGlo luciferase kit (Promega E2920) according to the manufacturer’s instructions. Relative light units were measured with BioTek Synergy 2 or Synergy LX plate reader (Agilent) using BioTek Gen5 software. Firefly luciferase values were normalized to renilla luciferase, with values set relative to a negative control value.

For MP10 reporter assays, C2C12s were seeded at 50, 000 cells/well in a 24 well plate. The following day, cells were transfected cells for 6 hours as described above. The next day, stimulations started using 10μM FSK, 18μM IBMX, 1, 5, and 10μM MP10 or DMSO control and continued for 6 hours until cell lysate was harvested in 1X passive lysis buffer, after which samples were processed as described above.

### Western Blot

For stimulation experiments, C2C12s were treated with cAMP agonists and then protein lysates were harvested for western blot. For CRISPRa western blots, cells were seeded at 10, 000 cells/well in a 6 well plate. Starting the next day, cells were stimulated for 6 days with 1 μg/ml Dox with media changes every other day. Cells were then harvested in RIPA buffer supplemented with protease inhibitor (Sigma 11836153001). Lysate was sonicated and cleared by centrifugation at 20, 000xg for 15 minutes, and protein was quantified using a BCA protein assay (ThermoFisher 23225). Protein (50μg/lane) was resolved by SDS-PAGE on a 7.5% gel (Bio-Rad 4561023) at 110V for ∼1 hour in tris-glycine SDS buffer. Proteins were transferred from the gel onto a nitrocellulose membrane using wet transfer at 110V for ∼1 hour in 20% methanol tris-glycine transfer buffer. The membrane was blocked in a solution of PBS-T with 0.5% powdered milk for 1 hour before an overnight incubation at 4℃ in primary antibody diluted in blocking solution (anti-Pde10a, AbCam 22782, 1:1000; anti-HA, Invitrogen PI26183, 1:2000; anti-beta-actin, Santa Cruz Biotechnology sc-47778, 1:1000). The next day, the membrane was washed three times (10 min each) with PBS-T and then incubated with a 1:4000 dilution of HRP-conjugated secondary antibody in blocking solution for 1 hour at room temperature. Membranes were washed three times with PBS-T (10 min each) and secondary antibody was detected using enhanced chemiluminescence substrate (ThermoFisher 32106), and membranes were imaged the membrane using a BioRad ChemiDoc and ImageLab software. For reprobing the membrane after imaging, the membrane was twice rinsed in a stripping buffer, washed twice with PBS-T, and then reblocked in blocking solution. Band intensity was quantified by densitometry using Fiji/ImageJ software. Briefly, rectangular lanes were drawn around each of the bands and profile plots were generated to calculate area under the curve for each band, which was normalized to the loading control and then set relative to the unstimulated group.

### Quantitative proteomics

For proteomics we used CRISPRa C2C12 cells described above, which expressed dox-inducible dCas9-VP64 and either a negative control LacZ gRNA or gRNA targeting the mouse sequence homologous to the *LINC00473* promoter (sg*Pde10a*). We seeded 2 biological replicates per condition in 15 cm plates and these cells treated with doxycycline for 4 d prior to extraction of protein for proteomics. Cell lysates were sonicated using a Branson SLPe Digital Sonifier set to 20% amplitude in a continuous mode, applying three 20-second cycles with one-minute resting on ice in between. The lysate was cleared by centrifugation at 20, 000 × g for 15 min. Protein concentration was determined using a BCA assay (Pierce 23225). Samples were then reduced with 5 mM dithiothreitol for 30 minutes at room temperature and alkylated with 12 mM iodoacetamide for 30 minutes in the dark. A 25 µg aliquot of each reduced and alkylated sample was processed and digested using single-pot, solid-phase-enhanced sample preparation (SP3) protocol (37). Briefly, a 10:1 bead-to-protein reaction was set up using SpeedBead Magnetic Carboxylate beads, rinsed with 80% ethanol, and digested with trypsin in 200 mM EPPS buffer (pH 8.2).

Following the release from SP3 beads, peptides were adjusted to 30% acetonitrile and labeled with 5 µL TMTpro 16-plex reagents (12.5 µg/µL in anhydrous acetonitrile) for 1 hour at room temperature. Reactions were then quenched with 5% hydroxylamine. A ratio check was performed using StageTip (38) cleanup and LC-MS2 analysis (39) of 5 µL combined from each TMT labeled channel and formed the basis for equal mixing. Following the analysis, each TMT channel signal was normalized and volumes for multiplexing were calculated, then pooled. The full multiplexed sample was desalted using a 100 mg tC18 Sep-Pak cartridge, eluted with 50% acetonitrile/1% formic acid then 70% acetonitrile/1% formic acid, and dried in a vacuum concentrator overnight.

The multiplexed peptide sample was reconstituted in 5% acetonitrile with 10 mM ammonium formate/ pH 8, briefly sonicated with two 12-second cycles, and centrifuged at 20, 000 × g for 5 minutes. Peptides were fractionated on an Accucore Vanquish C18+ column using a linear gradient from 0–40% solvent B (90% acetonitrile, 10 mM ammonium formate) over 18 minutes at 0.4 mL/min, followed by a wash at 100% B and re-equilibration. Ninety-six fractions were collected and combined into 12 pooled peptide fractions, which were desalted using StageTips. LC-MS/MS analysis (39) was performed on an Orbitrap Fusion Lumos mass spectrometer coupled to a Proxeon EASY-nLC 1200 LC pump system. Separation used a 90-minute gradient, ramping from 0 to 25% acetonitrile/0.1% formic acid over 82 minutes at 575 nL/min. MS1 scans (400–1600 m/z) were collected with FAIMS using three compensation voltages (−40, −60, −80), at 60, 000 resolution, 50 ms maximum injection time, and AGC target of 4e5. For each CV, MS2 scans were acquired with HCD fragmentation at 36%, 50, 000 resolution, and a 0.5 m/z isolation window.

Spectra were searched using MSFragger (40) in the FragPipe environment with the TMT16 workflow. Mass tolerances were set to ±10 ppm for precursors and 0.02 Da for fragments. Trypsin was set as proteolytic enzyme, allowing up to two missed cleavages. TMTpro fixed modifications included +304.2071 Da on peptide N-termini and lysine residues, and +57.0215 Da on cysteines. Methionine oxidation (+15.9949 Da) was included as a variable modification. The UniProt mouse proteome (accessed March 2025) served as the search database. False discovery rates for peptides and proteins were set to <1% using Philosopher (41). Quantitative data were processed in RStudio and Excel, and protein data from each TMT channel were normalized by a correction factor equalizing the summed intensity for all proteins in the channel. Significantly different proteins were revealed using a ttest implemented with the genefilter R package and p-values were adjusted using Benjamini-Hochberg multiple hypothesis correction. Significance was defined as Benjamini-Hochberg-corrected p-values <0.05 and log2 fold change <-0.6 or > 0.6.

### Neuronal differentiation

E14TG2a mouse ESCs were obtained from ATCC and were grown in high glucose DMEM supplemented with 15% Knockout Serum Replacement (Invitrogen), 1x non-essential amino acids (Cytiva), 1x sodium pyruvate (Cytiva), 4mM glutamine (Cytiva), 0.055 mM beta-mercaptoethanol (Gibco), 1% penicillin/streptomycin (Cytiva), and 1000 u/ml recombinant mouse leukemia inhibitory factor (R&D Systems or BioBasic), maintained at 37°C and 5% CO_2_. For passaging, ESCs were dissociated to single cells with Accutase and then seeded on plates coated with 0.2% gelatin (Sigma) (100, 000 cells per well of a 6 well plate), with media changes every day until cells were passaged again 2-3 days later. Mouse ESCs were differentiated to neurons using either an adherent differentiation protocol or an embryoid body-based protocol that specifically generates cortical neurons. For adherent differentiations, on day −4, we seeded 2.2×10^4^ cells/cm^2^ in N2B27 media (1:1 ratio of DMEF-12 (Cytiva, SH30023.01) and neurobasal (Gibco, 21103-049) with 1X N2 (Gibco, 17502-048), 0.5X B27 (Gibco, 17504-044), 1X glutamine (), 1X pen/strep (HyClone, SV30010), 1.82X β-mercaptoethanol (Gibco, 21985-023), 0.0025% BSA (Sigma-Aldrich, A7906-100G)) on 0.2% gelatin (Sigma-Aldrich, G1890-100G) coated plates. On day 0, lift cells with accutase (Millipore, SCR005) for 5 minutes at room temperature. Pellet cells by centrifuging at 120xg for 5 minutes. Resuspend cells in N2B27 media supplemented with 20ng/mL FGF2 (PeproTech, 100-18B-100UG) and seed 3.9×10^5^ cells/cm^2^ on Matrigel coated plates. On day 1, change media to N2B27 media. On day 2, 4 and 7, change N2B27 media. Cortical neurons (42) were generated using published procedures (43) with minor modifications. Upon dissociation of embryoid bodies, neurons were seeded on plates coated with Matrigel (Corning) or poly-L-ornithine (Sigma) and laminin (Sigma) as described (32), and were allowed to mature in neuronal media consisting of Neurobasal (Gibco) supplemented with 1x B27 for 8 days (with half media changes every other day). On day 8, neurons were fixed for immunocytochemistry or stimulated with 10μM FSK to activate cAMP signaling for gene expression analyses.

The human induced pluripotent stem cells line PGPC3_75, was derived from a healthy adult male donor, and was confirmed by whole genome sequencing to have no genetic variants known to affect neuronal function (31). iPSCs were grown in mTeSR Plus (STEMCELL Technologies 100-0276) and passaged using ReLeSR (STEMCELL Technologies 100-0483) as described (32, 44, 45). For conversion to neurons, iPSCs were transfected with PB-tetO-NGN2 (Addgene plasmid 197090) (46) followed by selection with 1 µg/mL puromycin. Puro resistant cells were seeded on Matrigel (Corning) or Cultrex (Biotechne) (1:60 to 1:100) and differentiated as described (45–47). After neurons were transitioned to neural differentiation medium, they were given half media changes every 3-4 days. On the day before stimulation, media was changed to neuronal medium without BDNF or GDNF. One day later, the cells were treated with 10 µM FSK/ 18 M µIBMX for 0, 1, 3, or 6 h, followed by harvesting RNA as described.

### Statistical Analysis

Statistical analysis was performed on GraphPad Prism 9.5.1. Data was assessed for normality using Shapiro-Wilk tests. If data met the assumptions of parametric tests, comparisons were analyzed by T-Test or ANOVA with Tukey or Dunnett multiple comparisons tests. If data failed to meet the assumptions of parametric tests, comparisons were analyzed by Mann-Whitney U or Kruskal Wallis tests with Dunn’s multiple comparisons test. Differences were considered statistically significant if p values were below 0.05. For information on statistical analysis of proteomics data, see above (*Quantitative proteomics*).

## RESULTS

### The LINC00473 promoter is functionally conserved in mice

Despite the absence of *LINC00473* exons, the *LINC00473* promoter is conserved in the mouse (13), including 2 conserved CREB binding sites (Supplementary Fig. S1 A and B), although it is unclear whether the mouse promoter is CREB-responsive like the *LINC00473* promoter. In the mouse, this sequence is a bona fide transcription start site (annotated in the FANTOM5 dataset as p3@Pde10a (48)) and is immediately upstream of a long *Pde10a* transcript variant that is expressed in rodents but is likely not conserved in humans (13) (Fig. 1A). To test the regulatory activity of the mouse homolog of the *LINC00473* promoter (herein called mouse *Pde10a p3*), we cloned this sequence and the human *LINC00473* promoters upstream of luciferase. The resultant reporter plasmids were transfected into 293T cells, which were then treated with increasing doses of the adenylyl cyclase agonist forskolin (FSK) to stimulate cAMP synthesis. Both the human *LINC00473* promoter and its mouse homolog supported dose-dependent increases in luciferase expression in response to FSK, peaking at 5-fold induction compared to unstimulated cells at the highest dose tested (Fig. 1B). Notably, the negative control reporter regulated by the *MMP1* promoter did not response to FSK (Fig. 1B). These results confirm that the mouse *LINC00473* homologe *Pde10a p3* is a cAMP-inducible promoter in reporter assays.

If mouse *Pde10a p3* is cAMP-inducible and regulates *Pde10a* as predicted (Fig. 1A and Supplementary Fig. S1), then cAMP stimulation should increase expression of the endogenous *Pde10a* gene. To test this prediction, we stimulated mouse C2C12 myoblasts with FSK and the broad-spectrum phosphodiesterase inhibitor IBMX to activate cAMP signaling, leading to phosphorylation of CREB (Fig. 1C) and activation of CREB-dependent gene expression (Fig. 1D). The negative control gene *Gapdh* did not respond to FSK/IBMX, and the positive control gene *Fos* was rapidly induced (Fig. 1D) as expected (49). In C2C12 cells stimulated with FSK/IBMX, we detected a pronounced induction of endogenous *Pde10a* transcript by ∼10-fold after 3 hours of stimulation, similar to the expression kinetics of human *LINC00473* (19) (Fig. 1D). FSK/IBMX-induced activation of mouse *Pde10a* was abrogated by overexpression of the dominant-negative CREB inhibitor ACREB (50), confirming that *Pde10a* is a bona fide CREB target gene in rodents (51). Our finding that mouse *Pde10a* expression is increased by cAMP signaling is consistent with previous studies of rodent pineal gland (52) and in rat INS-1 cells (51). These findings show that expression of mouse *Pde10a* is increased in response to cAMP signaling.

### The mouse homolog of the LINC00473 promoter regulates expression of Pde10a

Together, our results indicate that the cAMP-inducible human *LINC00473* promoter is functionally conserved in mouse (Fig. 1B) and that the mouse *Pde10a* gene downstream of that promoter is activated in response to cAMP signaling (Fig. 1D). However, it remains to be seen whether mouse *Pde10a p3* is responsible for cAMP-inducibility of the mouse *Pde10a* gene.

To directly test if *Pde10a p3* directly regulates *Pde10a* expression, we used CRISPRa with a gRNA targeting the promoter (Fig. 2A). C2C12 cells were infected with lentiviral vectors encoding the *Pde10a p3* gRNA or a negative control RNA targeting the prokaryotic *LacZ* gene (34) along with a lentiviral vector expressing a dox-inducible dCas9-VP64 fusion protein (36). Immunocytochemistry and qPCR confirmed dox-inducibility of dCas-VP64 (Fig. 2 B and C). For both the *LacZ* gRNA- and *Pde10a* gRNA-expressing CRISPRa C2C12 cells, there was no change in expression of the control gene *Gapdh*, with or without dox treatment (Fig. 2C). There was also no difference in dox-induced expression of dCas9-VP64 between the two gRNAs (Fig. 2C). Having confirmed efficacy and specificity of CRISPRa, we next examined expression of *Pde10a* using qPCR primers that annealed in exons of the *Pde10a* open reading frame (Fig. 2A). In the absence of dox, baseline *Pde10a* expression trended toward doubling in cells with the *Pde10a* gRNA compared to the *LacZ* gRNAs, suggesting leaky dox-independent expression of dCas9-VP64 (53). Despite this leaky activation of CRISPRa, dox treatment resulted in a 4-fold increase in *Pde10a* expression in cells expressing the *Pde10a* gRNA compared to cells expressing the *LacZ* gRNA (Fig. 2C). Together, these results suggest that activation of the mouse *Pde10a p3* promoter is sufficient to upregulate expression of *Pde10a* in the mouse.

**Figure 2.**
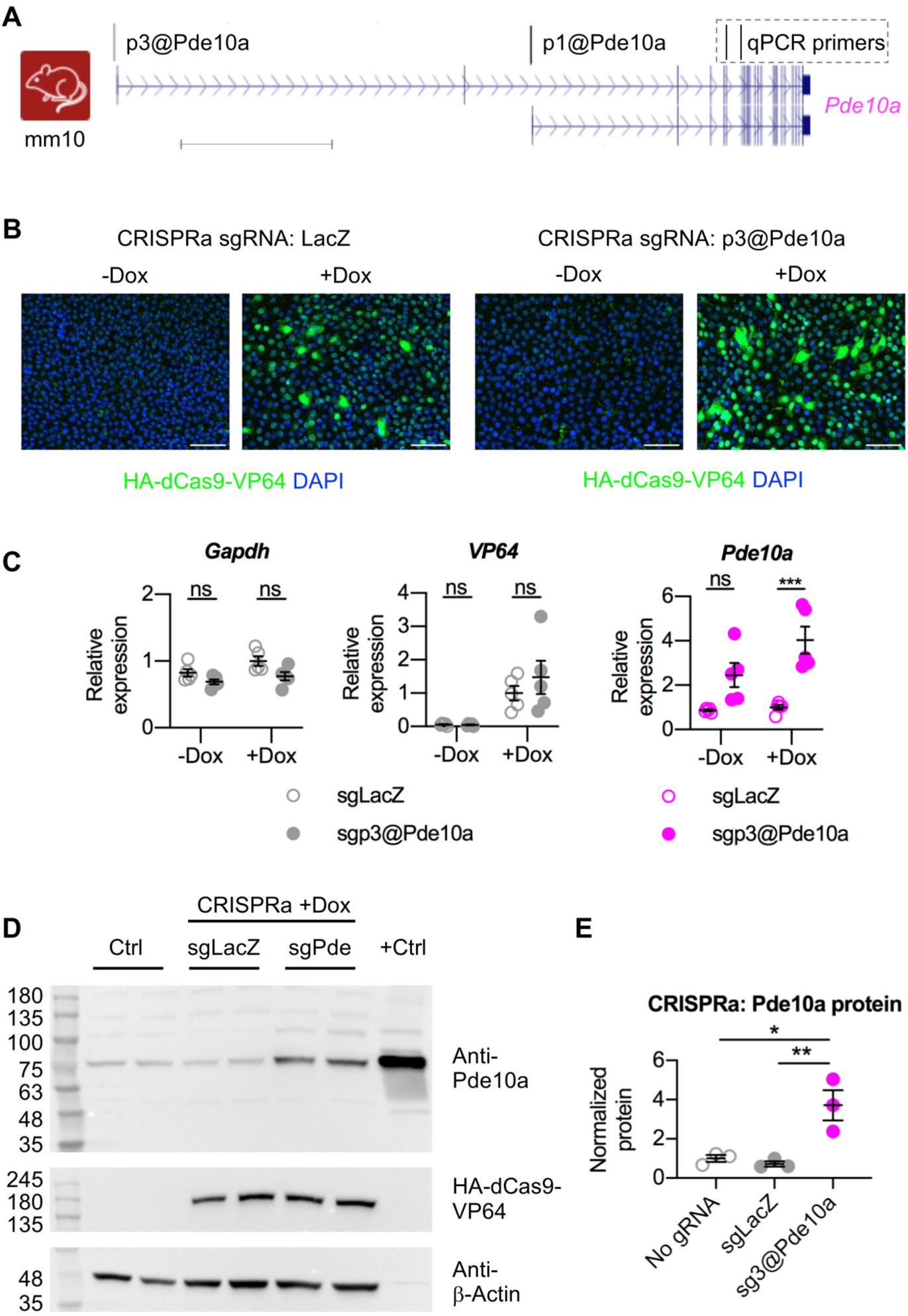
CRISPRa to test sufficiency of the conserved mouse promoter to regulate the endogenous *Pde10a* gene. (**A**) Schematic of mouse Pde10 locus showing the locations of the conserved promoter and the CRISPRa gRNA targeting the p3@Pde10a transcription start site (GRCm38/mm10). (**B**) Immunocytochemistry with to detect Dox-inducible HA-dCas9-VP64 in C2C12 cells with or without 1μg/mL Dox for 2 days. Scale bars = 200μm. (**C**) qPCR gene expression analysis in CRISPRa C2C12 cells with or without 1μg/mL Dox for 6 days, normalized to reference genes *Csnk2a2*/*Rer1*. Dots show individual data points, lines show mean, and error bars show standard error of the mean. (**D**) Representative western blots with the indicated antibodies in lysate from CRISPRa C2C12 cells (showing results from 2 biological replicates per condition) with brain lysate as a positive control for anti-Pde10a. (**E**) Densitometry analysis of western blot data for CRISPRa C2C12 cells (n=3 biological replicates per condition). (**C** and **E**) Data analyzed using one-way ANOVA with Tukey’s multiple comparisons test (*, P≤0.05; **, P≤0.01; ***, P≤0.001).

Having established that the mouse *Pde10a p3* promoter is cAMP-responsive and upregulates *Pde10a* transcript, we sought to test whether cAMP-inducible *Pde10a* could represent a form of negative feedback regulation, as has been described for other PDEs like *PDE4B* (54). However, such a feedback loop would require that transcriptional activation of *Pde10a p3* lead to a corresponding increase in Pde10a protein. To test this, we used CRISPRa to activate the cAMP-inducible promoter upstream of mouse *Pde10a* gene and then examined Pde10a protein levels by western blot. Due to suspected leakiness of dox-inducibility (Fig. 2C), we used only dox-treated cells for this analysis and compared the *Pde10a* gRNA to the negative control *LacZ* gRNA. Using this approach, we found that CRISPRa with the *Pde10a* gRNA led to a 3-fold increase in Pde10a protein levels compared to untreated C2C12 cells or CRISPRa cells expressing the *LacZ* gRNA (Fig. 2D and E). Together, these experiments show that activation of the mouse *Pde10a p3* promoter leads to increases in both *Pde10a* transcript and protein levels.

### Pde10a promotes differentiation of C2C12 cells

Having established that the human *LINC00473* promoter is functionally conserved in non-primates, we next sought to examine the function of the conserved sequence in species that lack a *LINC00473* gene. To this end, we used CRISPR activation (CRISPRa) to activate *Pde10a p3* in the myoblast-like C2C12 cells (55) followed by quantitative proteomics. C2C12 cells express muscle stem/progenitor cell markers like *Myod1* and *Pax7* (56), and either proliferate in the presence of growth factors or differentiate to muscle in low serum conditions (where they are also, to a lesser extent, able to differentiate to other mesenchymal tissues like bone, cartilage, and fat (57)). CREB has known roles in muscle stem/progenitor proliferation (58, 59) and differentiation (60), so C2C12s provide a tractable model to test cellular functions of CREB-induced *Pde10a* expression, which may provide insight into the consequences of *LINC00473* acquisition in primates.

To test functions of CREB-inducible *Pde10a* in the mouse, we compared proteomes of C2C12 CRISPRa cells that expressed a gRNA targeting the *Pde10a p3* or *LacZ*. This analysis revealed 459 differentially regulated proteins (log_2_ fold-change threshold: +/-0.6: Benjamini-Hochberg adjusted *p* value <0.05), of which 285 were upregulated and 174 were downregulated (Fig. 3A). Gene ontology enrichment analysis revealed that downregulated proteins had roles in purine deoxyribonucleotide metabolism and kinase signaling, whereas upregulated proteins were involved in cellular differentiation and mitochondrial electron transport (Fig. 3B). Analysis of select proteins confirmed upregulation of Pde10a, as expected, along with evidence of differentiation into muscle (Mylk2), bone (Tmem119), and cartilage (Sox9) (Fig. 3C). Downregulated proteins included the muscle stem cell marker Pax7, proteins known to support C2C12 proliferation (Pitx2 and its target Ccnd2 (61)), and the CREB target Mpp7 (58), which is essential for self-renewal of muscle stem cells *in vivo* (Fig. 3C). Consistent with these results, we found that CRISPRa-mediated activation of the CREB-dependent Pde10a promoter led to reduced C2C12 cell numbers (Supplementary Fig. 2A). To independently assess to role of CREB in the proliferation of C2C12 cells we overexpressed 2 different dominant negative CREB constructs, both resulting in decreased cell numbers, similar to *Pde10a* CRISPRa cells (Supplementary Fig. 2B). We also performed qPCR to test whether CRISPRa led to changes in muscle differentiation. We detected 4-fold upregulation of the muscle differentiation marker *Myog*, although notably this was abrogated by pharmacological inhibition of Pde10a with MP-10 (25, 62) (Supplementary Fig. 2D). Together, these results suggest that forced activation of the mouse *Pde10a p3* promoter in C2C12 cells leads to decreased cell numbers, presumably due to spontaneous differentiation.

**Figure 3.**
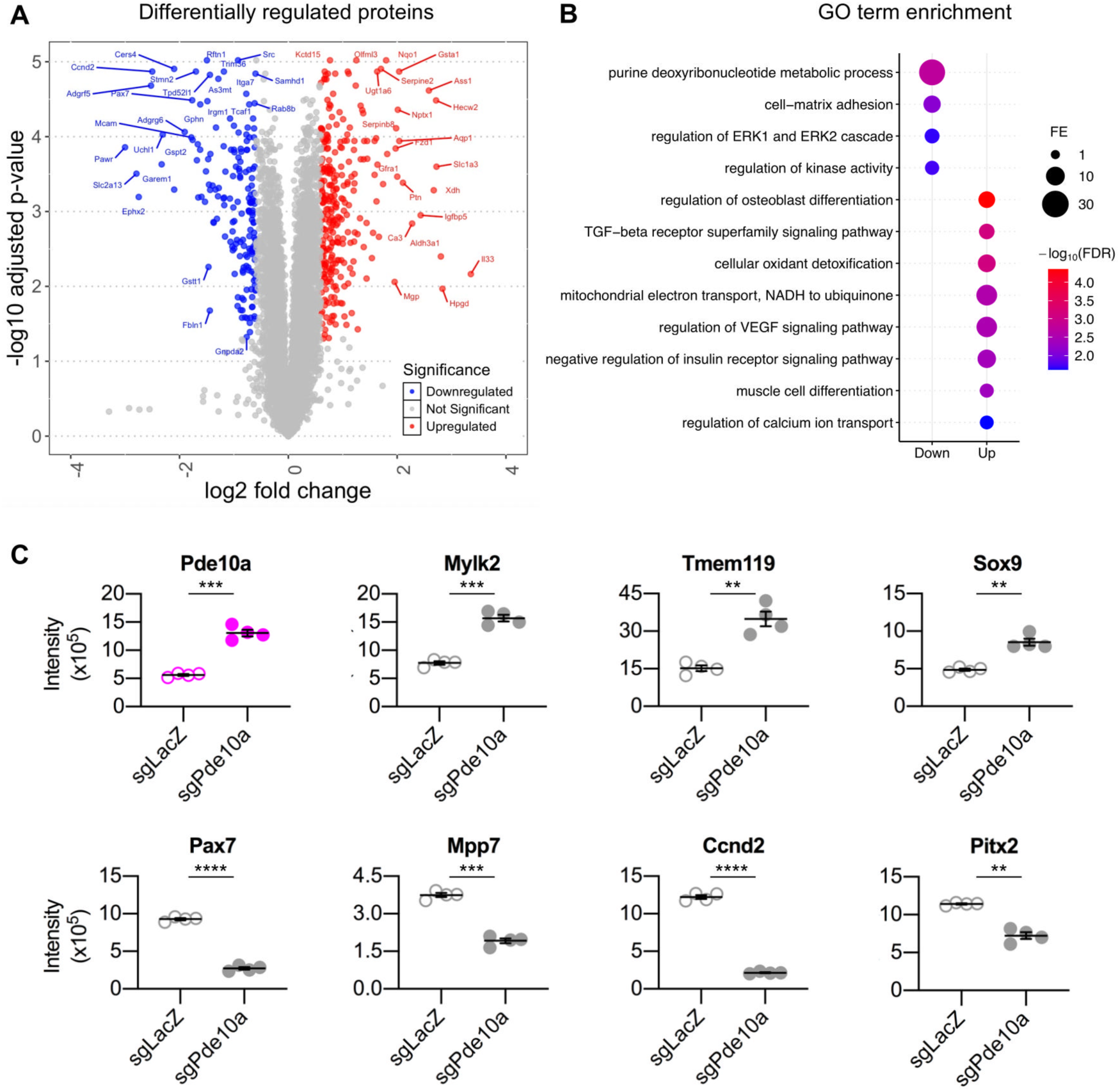
Quantitative proteomics analysis CRISPRa C2C12 cells overexpressing Pde10a from the mouse homolog of the *LINC00473* promoter. (**A**) Volcano plot displaying differentially regulated proteins in cells expressing *Pde10a* gRNA compared to cells expressing a negative control gRNA targeting *LacZ*. Red data points are upregulated in sgPde cells (log2 fold change >0.6), and blue data points are downregulated in sgPde cells (log2 fold change (<-0.6). Significance was assessed based on Benjamini-Hochberg-adjusted *p*-values <0.05. (**B**) Schematic depicting results of Gene Ontology (GO) term enrichment analysis of misregulated proteins. (**C**) Dot plots depicting abundance of proteins-of-interest from the proteomics analysis. Graphs display individual data points along with mean and standard error (n=4). Data analyzed using t-tests with Benjamini-Hochberg-adjusted *p*-values. multiple comparisons test (**, p<0.01; ***, *p*<0.001; \*\*\*\**p*<0.001). (**A**, **B**, **C**) All analyses include 2 replicates of 2 biological replicates per condition. (A, (C)

### Pde10a suppresses cAMP-induced gene expression

Having established that mouse *Pde10a* expression is induced by a cAMP-regulated promoter, we tested whether the resultant *Pde10a* protein acts as a negative regulator of cAMP signaling, as predicted based on its biochemical PDE activity (25). FSK stimulation activates adenylyl cyclase activity to drive cAMP synthesis, but this is countered by PDEs that then convert cAMP to AMP (Fig. 4A). PDEs, in turn, can be blocked by PDE inhibitors like IBMX to potentiate FSK-induced changes in cAMP-dependent gene expression (Fig. 4A). To test whether Pde10a specifically acts as a negative regulator of cAMP-induced gene expression, we tested whether the pharmacological Pde10a inhibitor MP-10 (25, 62) potentiated FSK-mediated activation of a reporter construct (Fig. 4A). We found that treatment with FSK alone drove a doubling in reporter expression, and this increased to a 4-fold induction when FSK was combined with IBMX (Fig. 4B). Notably, when combined with FSK, MP-10 treatment led to a dose-dependent potentiation of reporter expression, which was indistinguishable from effects of IBMX at the highest MP-10 dose (Fig. 4B). Our finding that inhibition of Pde10a activity leads to increased expression of a cAMP inducible reporter suggests that Pde10a could potentially act as a negative regulator of CREB.

**Figure 4.**
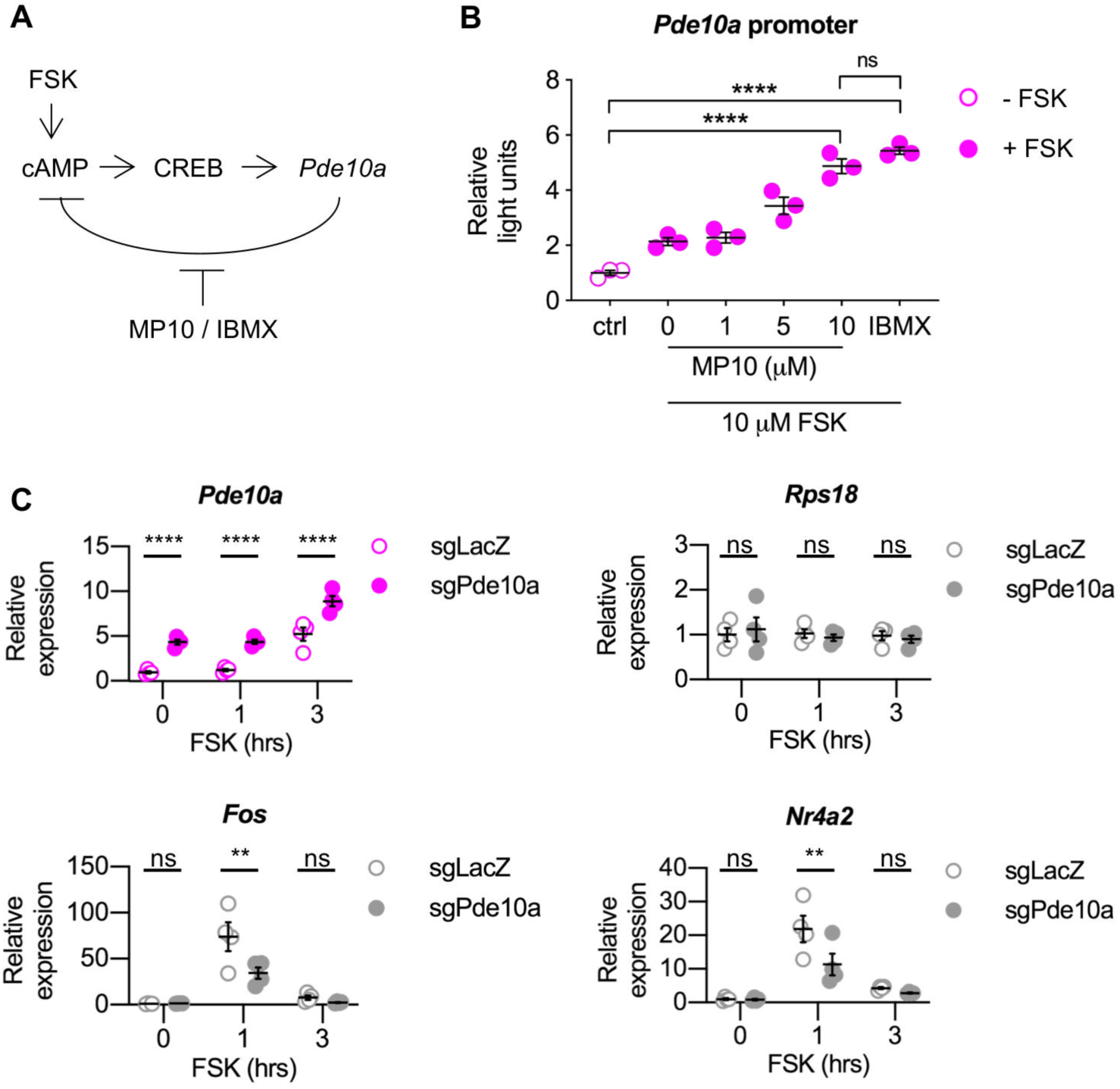
Testing role of Pde10a protein in regulation of cAMP-inducible gene expression. (**A**) Schematic representation of the cAMP signaling pathway highlighting targets of FSK, IBMX, and MP10. (**B**) Reporter assays in C2C12 cells transfected with a reporter plasmid where firefly luciferase is regulated by the mouse homolog of the *LINC00473* promoter. Cells were treated with 10μM FSK with or without phosphodiesterase inhibitors (1, 5 or 10μM MP10, or 18μM IBMX). Firefly luciferase was normalized to renilla luciferase control. Dots show individual data points (n=3 biological replicates), lines show mean, and error bars show standard error of the mean. Data analyzed using one-way ANOVA (*, P≤0.05; **, P<0.01; ****, P≤0.0001). (**C**) qPCR gene expression analysis in dox-treated CRISPRa C2C12 cells treated with or without FSK treatment Data are normalized to reference genes *Csnk2a2*/*Rer1*. Dots show individual data points (n=3-4 biological replicates, lines show mean, and error bars show standard error of the mean. Data analyzed using two-way ANOVA (**, P<0.01; ****, P≤0.0001).

To directly test the role of inducible *Pde10a* expression in CREB-dependent gene expression, we used CRISPRa to test whether elevating Pde10a protein via the mouse *Pde10a p3* promoter would affect FSK-induced gene expression in C2C12 cells. As expected, CRISPRa with sgPde10a led to a ∼5-fold upregulation of Pde10a compared to *LacZ* gRNA, and Pde10a expression was further increased in both conditions by 3h treatment with FSK (Fig. 4C). Notably, neither CRISPRa with sgPde10a nor FSK had any significant effect on the negative control gene Rps18 (Fig. 4C). We next tested expression on the CREB target genes *Fos* and *Nr4a2*. Both *Fos* and *Nr4a2* were unaffected by sgPde10a CRISPRa in unstimulated cells, whereas their induction after 1h of FSK stimulation was significantly ablated by 50% (Fig. 5C). These results demonstrate that forced activation of *Pde10a* expression via the *p3* promoter directly interferes with CREB-dependent gene expression.

**Figure 5.**
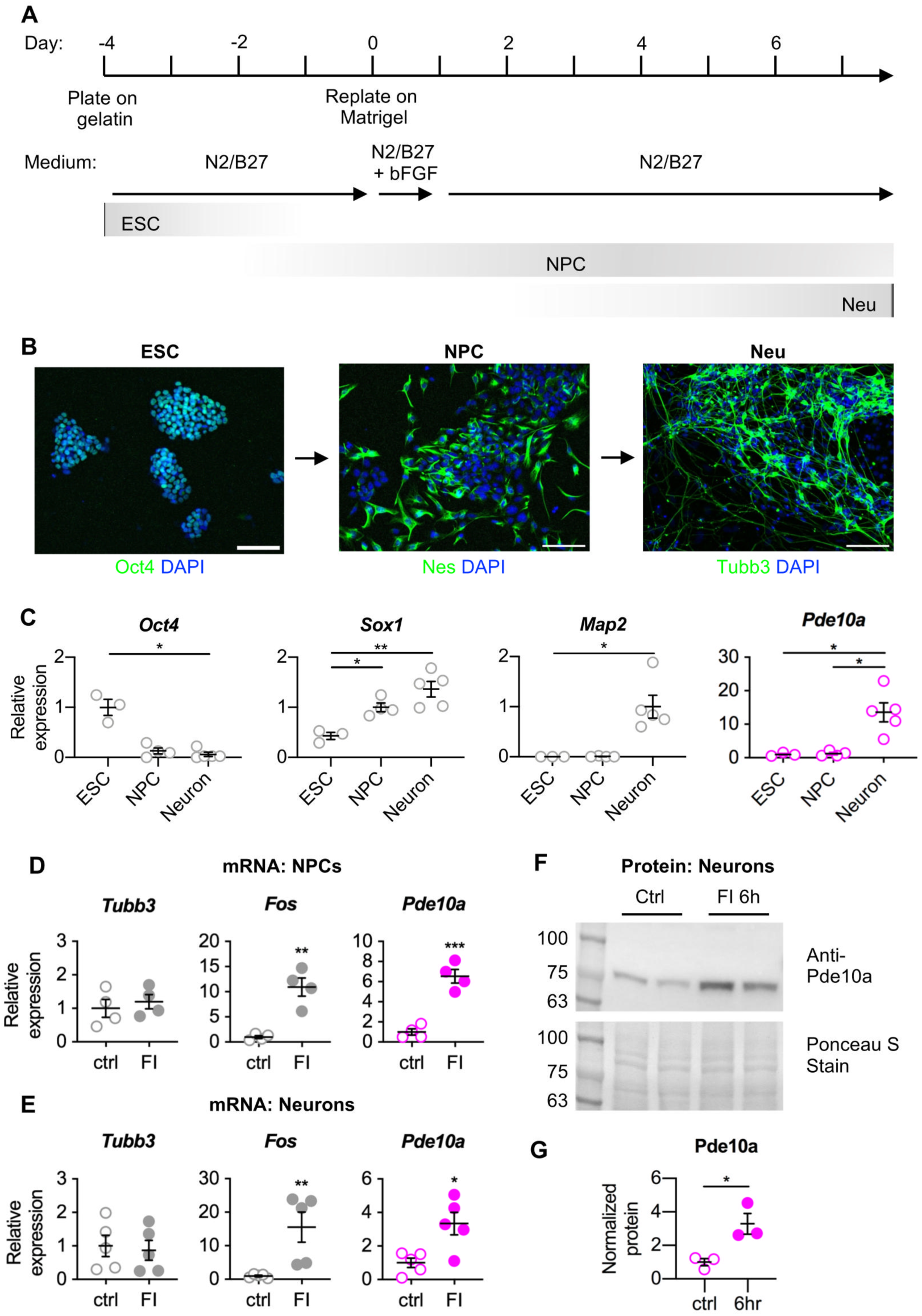
*Pde10a* is induced by cAMP signaling in mouse neurons derived from mESCs. (**A**) Schematic depicting adherent differentiation procedures used to differentiate mESC to neural progenitor cells (day 2) and neurons (day 7 and beyond). (**B**) Cell identity was visually assessed by immunocytochemistry with anti-Oct4 in undifferentiated mESCs, anti-Nestin at day 2 of the differentiation, and anti-Tubb3 at day 7, and all images include counterstain with DAPI. Scale bar = 200 micrometers. (**C-E**) Differentiations were also assessed by qPCR gene expression analysis in mESC, NPC, and neurons (C), and either NPCs (**D**) or neurons (**E**) that were treated with or without 3h stimulation with 10 µM FSK / µM 18 IBMX. qPCR data were normalized to reference genes *Gapdh* / *Rps18* (n= 3-5 biological replicates). (**F**) Representative western blots with the indicated antibodies in lysate from FSK/IBMX-stimulated mESC-derived neurons (showing results from 2 biological replicates per condition). (**E**) Densitometry analysis of western blot data for CRISPRa C2C12 cells (n=3 biological replicates per condition). (**C, D, E** and **G**) Data analyzed using unpaired two-tailed t-tests or Mann-Whitney test (*, P≤0.05; **, P≤0.01).

### Mouse Pde10a is cAMP-inducible in neurons

Having established that mouse *Pde10a* expression was increased by cAMP signaling in C2C12 cells, we next sought to determine whether cAMP-inducibility occurs in other mouse cell types. Given the importance of *Pde10a* in the brain (22, 25), we tested neuronal expression dynamics. This was particularly important in light of recent findings that synaptic activity in mouse neurons led to increased expression of *Pde10a* exons near the *p3* promoter without significant changes in downstream exons of the *Pde10a* open reading frame (13). To test whether cAMP signaling specifically activates *Pde10a* mRNA in mouse neurons, we differentiated mouse embryonic stem cells (ESC) to neural progenitor cells (NPC) and neurons (Neu) using an adherent differentiation protocol (42, 43) (Fig. 5A). We confirmed successful differentiations using immunocytochemistry. As expected, ESCs expressed the pluripotency marker Oct4, NPCs expressed the intermediate filament protein Nes, and Neurons expressed the terminal differentiation marker Tubb3 (beta-III-tubulin) (Fig. 5B). We also used qRT-PCR to test expression of Pde10a through the neural differentiation process, which revealed a ∼10-fold increase in Pde10a expression in neurons, coinciding with a pronounced increase in expression of the neural marker *Map2* (Fig. 6C).

**Figure 6.**
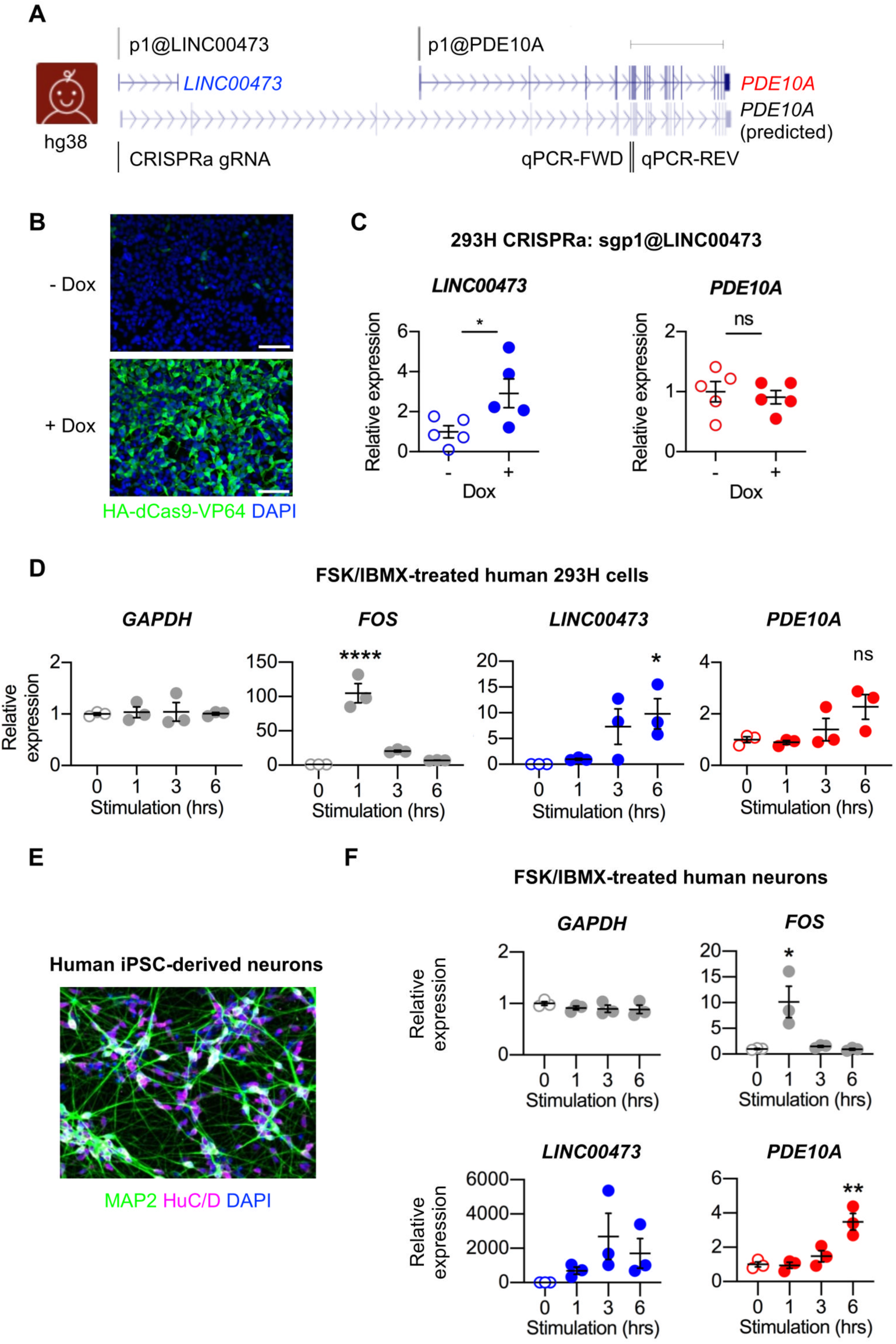
Differential regulation of *LINC00473* and *PDE10A* in human cells. (**A**) Schematic of human *PDE10A* and mouse *Pde10a* loci. Promoter labels correspond to FANTOM5 CAGE-seq transcriptions start sites (TSS). Schematic of the human *LINC00473* / *PDE10A* genomic locus depicts the location of the CRISPRa gRNA near sgp1@LINC00473, *PDE10A* qPCR primers, and a predicted *PDE10A* transcript starting near sgP1@LINC00473 (XM_011535387.4). Representative transcripts are displayed for each TSS / transcription unit. Modified from UCSC Genome Browser (GRCh38/hg38, GRCm38/mm10); scale bar = 100 kb. (**B**) Visualization of inducible HA-tagged dCas9 in CRISPRa 293H cells with or without 48 hour treatment with 2 µg/ml Dox. **(C)** qPCR gene expression analysis of *LINC00473* and *PDE10A* in CRISPRa 293H cells with or without 0.1 µg/mL Dox for 7 days (n = 5). Data were analyzed by unpaired two-tailed t-tests (* p ≤ 0.05). (**D**) qPCR gene expression analysis of *LINC00473* and *PDE10A* in FSK/IBMX-treated 293H cells (n = 3). Data were analyzed by one-way ANOVA with Dunnett multiple comparisons test (* p ≤ 0.05; ****, P<0.0001). (**E**) Visualization of neuronal markers MAP2 and HuC/D in human iPSC-derived neurons. (**F**) qPCR gene expression analysis of *LINC00473* and *PDE10A* in FSK/IBMX-treated iPSC-derived neurons (n = 3). Data were analyzed by one-way ANOVA with Dunnett multiple comparisons test (* p ≤ 0.05; ****, P<0.0001).

Having established a neural differentiation model, we next sought to test whether neural lineage cells induced expression of *Pde10a* in response to cAMP signaling induced by FSK / IMBX. As expected, the negative control gene *Tubb3* did not change in NPCs or neurons, and the positive control *Fos* was upregulated by cAMP in both cell types (Fig. 5 D and E). FSK/IBMX also led to *Pde10a* mRNA expression increasing by 5-fold and 3-fold after 3 h stimulation in NPCs and neurons, respectively (Fig. 5 D and E), and led to Pde10a protein levels increasing a 3-fold in neurons after 6 h (Fig. 5 F and G). To confirm cAMP-responsiveness of Pde10a in neurons in another model, we also differentiated mouse ESCs to cortical neurons using a widely-used embryoid body based-differentiation approach (Supplementary Fig. S3A) (42). As expected, the resultant neurons (Supplementary Fig. S3B) increased expression of *Pde10a* by 3-fold after 3 h stimulation with FSK / IBMX (Supplementary Fig. S3C). Similar to our results with mouse C2C12 cells, our findings with mouse ESC-derived neurons revealed that activation of cAMP signaling leads to increased expression of *Pde10a*.

### PDE10A is not induced by cAMP signaling in human cells

Our results thus far suggest that the mouse homolog of the *LINC00473* promoter is functionally conserved in mice, where it drives cAMP-inducible expression of *Pde10a*, which may then in turn act as a negative feedback regulator to dampen CREB-dependent gene expression. In human cells, this conserved CREB-inducible promoter drives expression of *LINC00473* (Fig. 6A), suggesting that *LINC00473* has hijacked this promoter in primates, where it presumably directs CREB-induced transcription of *LINC00473* instead of *PDE10A*. To test this hypothesis, we next assessed transcriptional consequences of activating the *LINC00473* promoter in 293H human cells using dox-inducible CRISPRa with a gRNA targeting the *LINC00473* (33, 36). After infection and selection, we confirmed by immunocytochemistry that most cells exhibited dox-inducible expression of dCas9-VP64 (Fig. 6B). We also found that treatment of *LINC00473* CRISPRa cells with dox resulted in a 2-fold increase in expression of *LINC00473* with no change in *PDE10A* (Fig. 6C), similar to previous CRISPRa studies in human adipocytes (63). These results suggest that the *LINC00473* promoter activates only *LINC00473*, and not its downstream neighbor *PDE10A*.

Having confirmed that *LINC00473* and *PDE10A* are independently regulated in human cells, we next tested whether *PDE10A* responds to cAMP signaling in human cells, which are known to upregulate *LINC00473* (13, 14, 17, 19, 63) in response to cAMP. As a first step, we tested the 293H cell line used for our *LINC00473* CRISPRa analysis. When stimulated with FSK/IBMX, 293H cells upregulated *FOS* at 1 h and *LINC00473* at later times, as expected (13, 19) (Fig 6D). Similar results were observed in MCF-7 cells, which upregulated *NR4A2* and *LINC00473* in response to FSK, but not *PDE10A* (Supplementary Fig. S6). These results suggest that *LINC00473* and *PDE10A* are independently regulated and human *PDE10A* is not induced by cAMP signaling in human cells.

Having determined that PDE10A is unresponsive to cAMP signaling in human cell lines, so we next tested whether *PDE10A* responds to cAMP signaling in human neurons. To this end, we used human induced pluripotent stem cells from a healthy adult donor (31) and directly converted these cells to neurons using *Ngn2* (46, 47, 64). Resultant neurons expressed MAP2 and HuC/D (Fig. 6E), and stimulation with FSK/IBMX led to upregulation of *FOS* with no change in *GAPDH* as expected (Fig. 6F). Analysis of *LINC00473* suggested upregulation with substantial ∼2, 000-fold change compared to baseline, although this trend did not meet statistical significance due to variability in the 3 biological replicates (Fig. 6F). Surprisingly, *PDE10A* was upregulated in response to FSK/IBMX in human neurons after 6 h, although the kinetics were delayed compared to mouse neurons, which significantly upregulated *Pde10a* after 3 h (Fig 6F). Our findings reveal that *PDE10A* is upregulated in human neurons after cAMP signaling, although with a notable delay of several hours compared to mouse neurons. Together with results of our CRISPRa experiments, these results reassert that human *LINC00473* and *PDE10A* are distinct, independently regulated genes, and suggest that acquisition of *LINC00473* in primates led to changes in the regulatory dynamics of *PDE10A* expression.

## DISCUSSION

This study provides unexpected insight into the evolutionary consequences of the newly evolved, primate-specific lncRNA *LINC00473*, which is induced by cAMP signaling and sits upstream of the *PDE10A* gene on the same chromosomal strand. *PDE10A* is a negative regulator of cAMP signaling and is unresponsive to cAMP in human cells. We confirmed that the *LINC00473* promoter is conserved in rodents and we found that this promoter supports cAMP inducible gene expression of both reporter constructs and in the context of the endogenous mouse *Pde10a* gene. CRISPRa also confirmed that activating the mouse sequence homologous to the *LINC00473* promoter is sufficient to drive transcription of *Pde10a*. We also found that inhibition of Pde10a enzyme activity increases expression of a cAMP inducible promoter. Together, these findings suggest that acquisition of *LINC00473* in primates split the ancestral Pde10a gene into 2 independently regulated transcription units: the cAMP inducible *LINC00473* and *PDE10A*, which is no longer responsive to cAMP.

lncRNAs are increasingly recognized for their ability to regulate expression of nearby and distal genes which can cause global changes to a cell (5–7). Here, we propose a gene regulatory function for *LINC00473* wherein it has altered the cAMP inducibility and expression of its neighbouring gene *PDE10A* independent of RNA sequence in processed *LINC00473* transcripts. *PDE10A* is widely conserved in many vertebrate species as a negative regulator of intracellular cAMP (25, 27, 65). We found that *PDE10A* is not cAMP inducible in human MCF7 cells and that changes in PDE10A mRNA levels are delayed in human neurons compared to mouse C2C12 cells and neurons (with similar trends in human 293H cells). This is consistent with findings in human iPSC-derived responding to synaptic stimulation (13). Conversely, we found that mouse *Pde10a* is activated by cAMP signaling in mouse C2C12 cells and mESC-derived neurons, suggesting that *Pde10a* may be inducible in non-primate species lacking *LINC00473*. We believe this change in inducibility is due to the evolution of the *LINC00473* gene, which has hijacked the cAMP inducible promoter once belonging to *Pde10a*. Since Pde10a enzymatic activity suppresses cAMP-dependent gene expression, we hypothesize that *LINC00473* has abrogated a cAMP-inducible negative feedback loop. Other PDEs like PDE4B have been shown to be regulated under cAMP feedback loops (54, 66), and our results show how regulation of these genes can change with the evolution of new lncRNA genes, leading to changes in downstream cAMP signaling.

Our findings provide data supporting alternative mechanisms by which new lncRNA genes may evolve. The birth of a new gene requires regulatory elements – including promoters, enhancers, splice sites, and poly-adenylation sequences – to enable transcription and processing of the transcript (12). Therefore, many newly evolved genes, including lncRNAs, arise from ancestral gene regulatory elements (9, 10). Some newly evolved lncRNAs arose from bidirectional promoters of protein coding genes or as a result of transcriptional initiation from enhancers (10, 67, 68). Our findings suggest that *LINC00473* acquired its regulatory elements by hijacking an inducible *Pde10a* promoter. Though there is some evidence of promoter hijacking events occurring in the acquisition of de novo protein coding genes (9), our results provide an example of promoter hijacking specifically disrupting a transcriptional feedback loop by redirecting activity of an alternative promoter for *Pde10a* while leaving its primary promoter (48) and transcript intact This ensures that baseline functions of *Pde10a* can continue unimpeded, while cAMP-inducible *Pde10a* is specifically disrupted or delayed.

CRISPRa was effective at increasing *Pde10a* transcript/protein levels in C2C12 cells and serves as the basis for future experiments involving alterations in *Pde10a* expression in these cells and primary myoblasts. Our pharmacological data revealed that inhibition of Pde10a alone was functionally comparable to the broad spectrum PDE inhibitor IBMX with respect to potentiating FSK-mediated induction of a CREB-regulated reporter gene. This finding suggests that Pde10a is a major source of PDE activity in these cells. cAMP signaling promotes muscle growth (59) and Pde10a inhibitors improved cell pathology in zebrafish models of Duchenne Muscular Dystrophy (65). Therefore, the increases in *Pde10a* transcription from CRISPRa may affect overall cell growth in vitro with C2C12 cells and primary myoblasts or in vivo in mouse models of muscle injury or degeneration. We detected some leakiness and only ∼2-3-fold induction with the Dox-inducible dCas9-VP64-based CRISPRa system, so future experiments will use the more robust constitutively expressed dCas9-VPR system (69). Additionally, the inverse experiments can be performed using CRISPR interference (CRISPRi) to decrease *Pde10a* expression to test if Pde10a is necessary for myoblast proliferation. CRISPRi works much the same as CRISPRa, but instead is fused to the transcriptional repressor KRAB. Together, these methods can be used as an adjunct to explore potential effects on growth related to *Pde10a* expression and cAMP regulation.

*PDE10A* is implicated in neuromuscular and psychiatric disorders affecting the communication between striatal medium spiny neurons (22, 25). PDE10A inhibitors were also tested as possible treatment options due to their ability to modify cAMP signaling (23). The efficacy of PDE10A inhibitors has been tested in animal models, like the mouse, where they have shown promise in their treatment. However, these same results do not seem to translate to humans(23, 70). This lack of therapeutic benefit in humans may be due to the difference in *PDE10A* regulation, wherein the human protein is no longer involved in cAMP regulating feedback loops that are still active in the mouse. This difference in *PDE10A* inducibility should be considered in future experiments when testing the efficacy of *PDE10A* inhibitors in mouse models. One potential approach would be to humanize the mouse *Pde10a* locus by disrupting communication between the cAMP-inducible promoter and the Pde10a open reading frame, perhaps by insertion of a premature polyadenylation site (13, 71), thereby preventing cAMP inducibility of *Pde10a*.

In summary, our findings demonstrate conservation of the promoter for the primate-specific gene *LINC00473* in both sequence and regulatory function within the mouse where it instead serves as a promoter for *Pde10a* to regulate cAMP inducible transcription of the gene. Our findings also suggest that *Pde10a* inducibility by cAMP in non-primate species may support a negative feedback loop to limit cAMP signaling. Overall, these results suggest that the acquisition of *LINC00473* in primates may have separated Pde10a from an ancestral cAMP-inducible promoter, and this may have led to distinct cAMP regulatory dynamics in primates and non-primate mammals.

## ACKNOWLEDGEMENTS

This work was supported by Discovery Grant (RGPIN-2017-05469) and Discovery Development Grant (DDG-2024-00025) from the Nature Science and Engineering Research Council (NSERC) to PJR; Internal and Startup Grants from the University of Prince Edward Island to PJR; Atlantic Cancer Research Grant from JD Irving Ltd and the Canadian Cancer Society and Canadian Cancer Society Emerging Scholars Grant to JPM; Jeanne and J.-Louis Lévesque Professorship to APJ; NIH/NIGMS R01 grant GM132129 and R35 grant GM156406 to JAP; John Evans Leadership Fund grant from the Canada Innovation Foundation with additional funding from the Jeanne and J.-Louis Lévesque Foundation and Innovation PEI to PJR and JPM. SJS was supported by a NSERC Canada Graduate Scholarship (CGS-M), NSERC Undergraduate Student Research Award (USRA) and a Summer Studentship Award from the Stem Cell Network. VSB, EC, and BIM were also supported by NSERC USRAs. We thank Dr. Rebecca Mok for helpful feedback on the manuscript, Drs. Chelsea Martin and Paola Marcato for experimental advice, Dr. James Ellis for kindly sharing plasmids and cell lines, and Drs. Stephen Scherer and James Ellis for kindly sharing human iPSC line PGPC3_75. We also thank Kaileigh Rowe, Clint Maramag, Marcus Gauthier, and Jason Ryan for technical assistance. We also thank Dr. Steven P. Gygi and the Taplin Mass Spectrometry Facility at Harvard Medical School for use of their mass spectrometer.

## AUTHOR CONTRIBUTIONS

Shelby J. Squires (Conceptualization, Investigation, Methodology, Formal analysis, Writing – original draft, Writing – review and editing, Visualization), Brandon S. Smith (Conceptualization, Investigation, Methodology, Formal analysis, Writing – original draft, Writing – review and editing, Visualization), Mukhayyo Sultonova (Methodology, Formal analysis), Emma J. Campbell (Methodology), V. Shruthi Bandi (Methodology), Brandon I. MacDonald (Methodology), Joao A. Paulo (Investigation; Methodology; Funding acquisition), Adam P. Johnston (Conceptualization, Methodology, Writing – review and editing, Funding acquisition), J. Patrick Murphy (Conceptualization, Formal analysis, Writing – review and editing, Supervision, Funding acquisition), and P. Joel Ross (Conceptualization, Investigation, Methodology, Formal analysis, Writing – original draft, Writing – review and editing, Visualization, Supervision, Funding acquisition).

---

**Figure S1.**
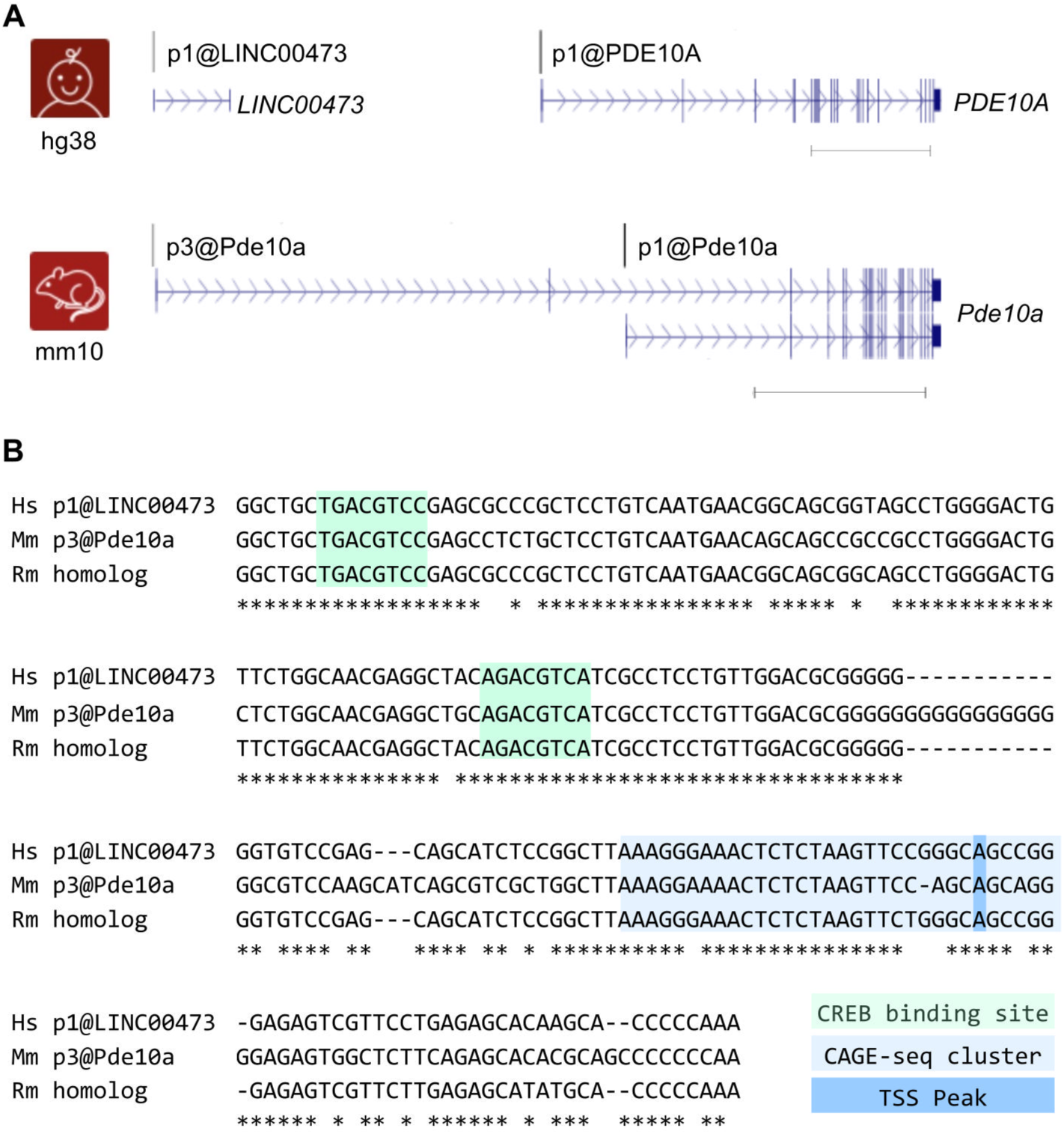
Conservation of the *LINC00473* promoter between humans and mice. (**A**) Schematic of *PDE10A/Pde10a* loci in the human and mouse genomes displaying locations of transcription start sites and transcription units. Promoter labels correspond to FANTOM5 CAGE-seq transcriptions start sites (TSS), with p1 representing the primary TSS for each gene (e.g., p1@LINC00473 for LINC00473). For each transcription unit, a single representative transcript is displayed (Human: NR_026861.1 for *LINC00473* and NM_001130690.3 for *PDE10A*; mouse: NM_001290707.1 and NM011866.2 for the long and short Pde10a transcripts respectively). Modified from UCSC Genome Browser (GRCh38/hg38, GRCm38/mm10); scale bar = 100 kb. (**B**) Alignment of the human *LINC00473* promoter (−160 to +40) with homologous sequences from the mouse (Mm) and Rhesus macaque (Rm) genomes. Asterisks indicate positions with identical sequences in all three genomes. Highlighted are putative CREB binding sites (green), CAGE-seq clusters (blue), and predicted peak TSS (dark blue) based on CAGE-seq data.

**Figure S2.**
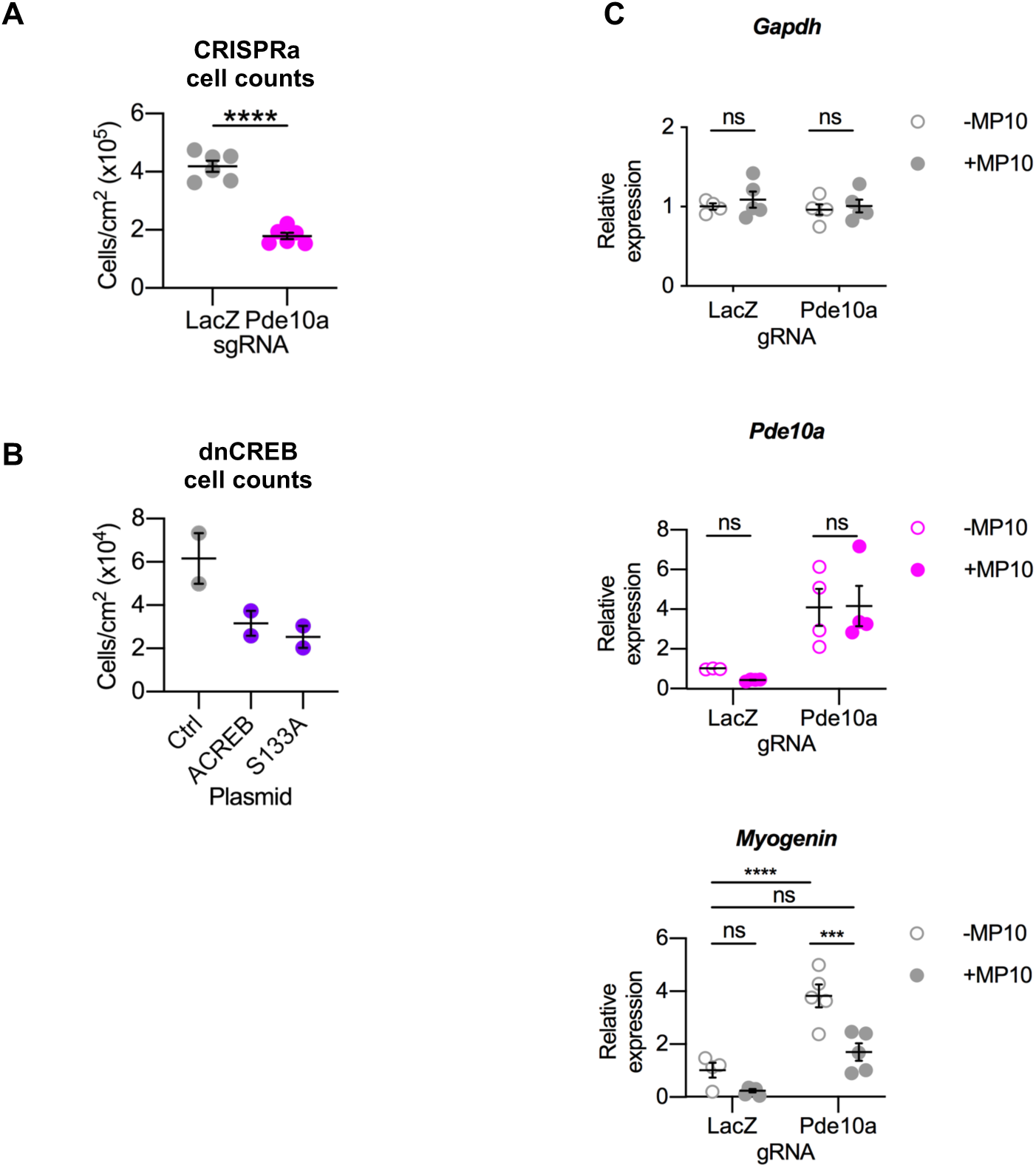
*Pde10a* upregulation promotes C2C12 differentiation. (**A**) Cell counts of CRISPRa C2C12 cells expressing gRNAs targeting either the mouse homolog of the *LINC00473* promoter (Pde10a) or a negative control *LacZ* gRNA (n=6 biological replicates), which were allowed to grow for 5 days (4 in dox) before counting. Data analyzed using unpaired two-tailed t-tests (****, *p*≤0.0001). (**B**) Cell counts of C2C12 cells that grew for 3 days while expressing dominant negative inhibitors of CREB, ACREB or S133A (n=2 biological replicates). (**C**) qPCR gene expression analysis in dox-treated CRISPRa C2C12 cells treated with or without the Pde10a inhibitor MP10, normalized to reference genes *Csnk2a2*/*Rer1*. Dots show individual data points (n=3-4 biological replicates, lines show mean, and error bars show standard error of the mean. Data analyzed using two-way ANOVA (***, P<0.001; ****, P≤0.0001).

**Figure S3.**
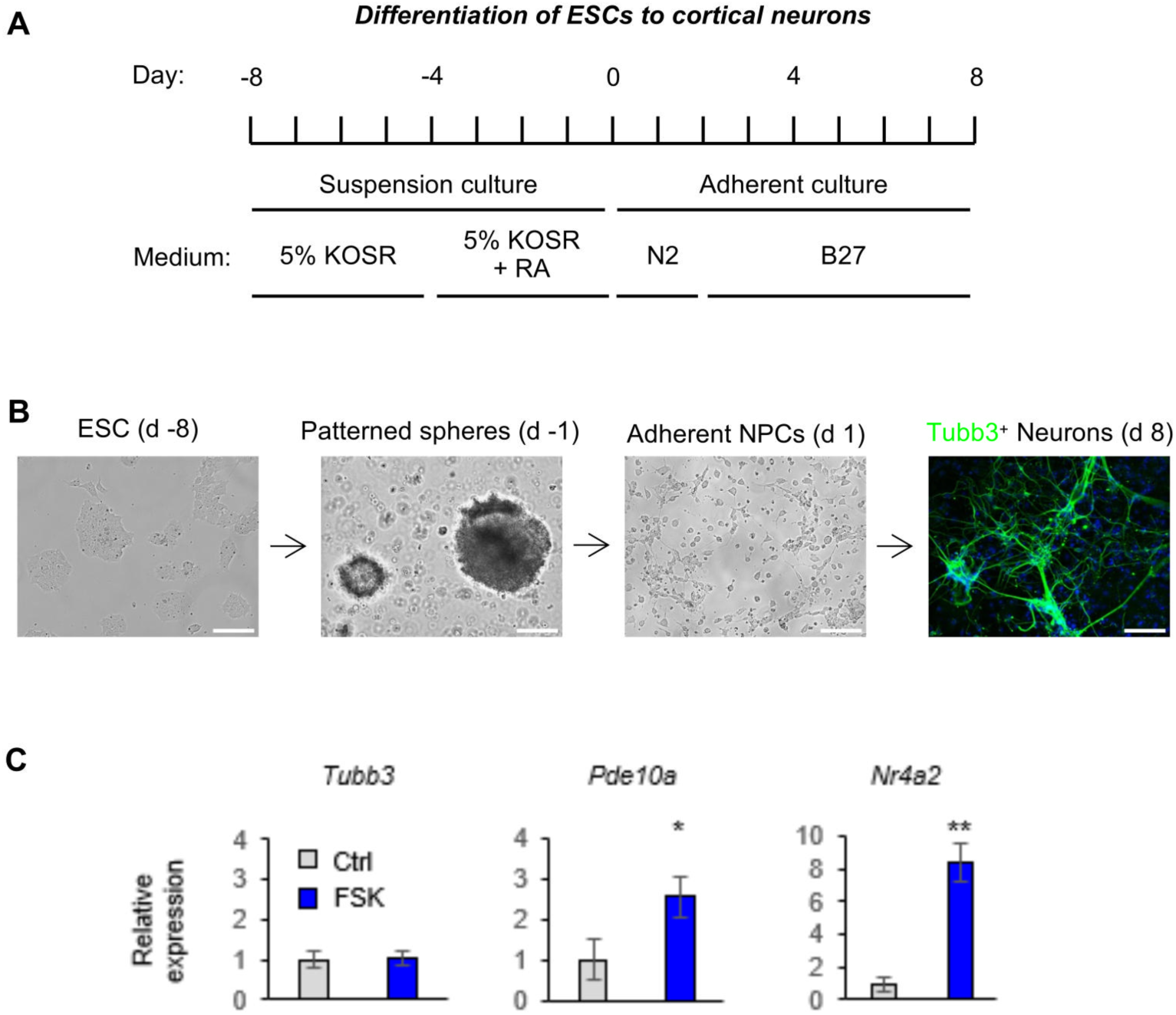
Pde10a is induced by cAMP signaling in mouse embryonic stem cell (mESC)-derived cortical neurons. (A) Schematic depicting the differentiation procedures used to differentiate mESC to neurons over a period of 16 days (d). (B) Visualization of mESC-derived neurons by immunocytochemistry with anti-beta-III-tubulin (Tuj1). Scale bar = 200 micrometers. (C) qPCR gene expression analysis in mESC-derived neurons with or without 3h stimulation with 10μM FSK normalized to reference genes *Gapdh* / *Rps18* (n = 3-4 biological). Graphs display mean and standard error. Data were analyzed by t-test (* p ≤ 0.05; **, P<0.01).

**Figure S4.**
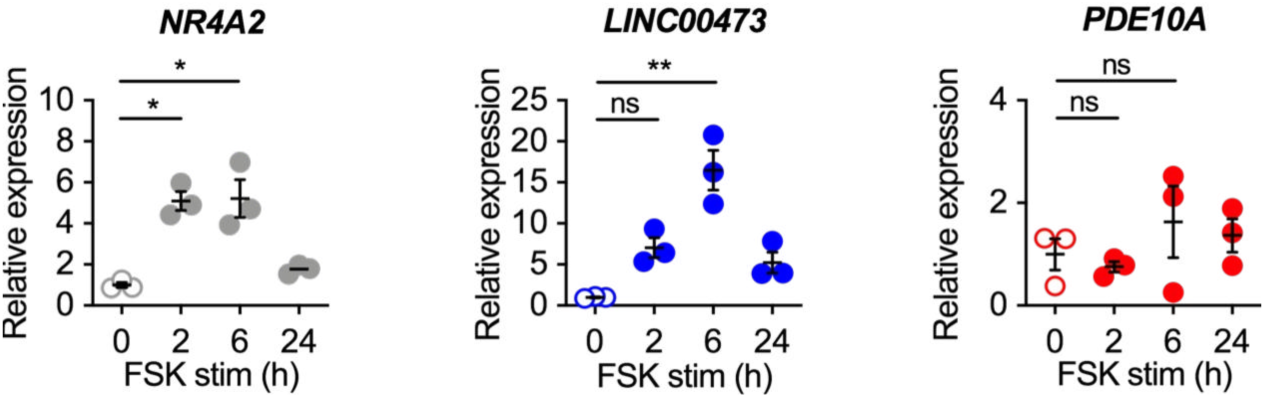
Differential regulation of *LINC00473* and *PDE10A* in human MCF-7 cells. qPCR gene expression analysis of the indicated genes in MCF-7 cells after stimulation with 50 uM FSK (n = 3 biological replicates). Dots show individual data points, lines show mean, and error bars show SEM. Data were analyzed by ANOVA with Dunnett posthoc test (* p ≤ 0.05; **, P<0.01).

**Table S1.**
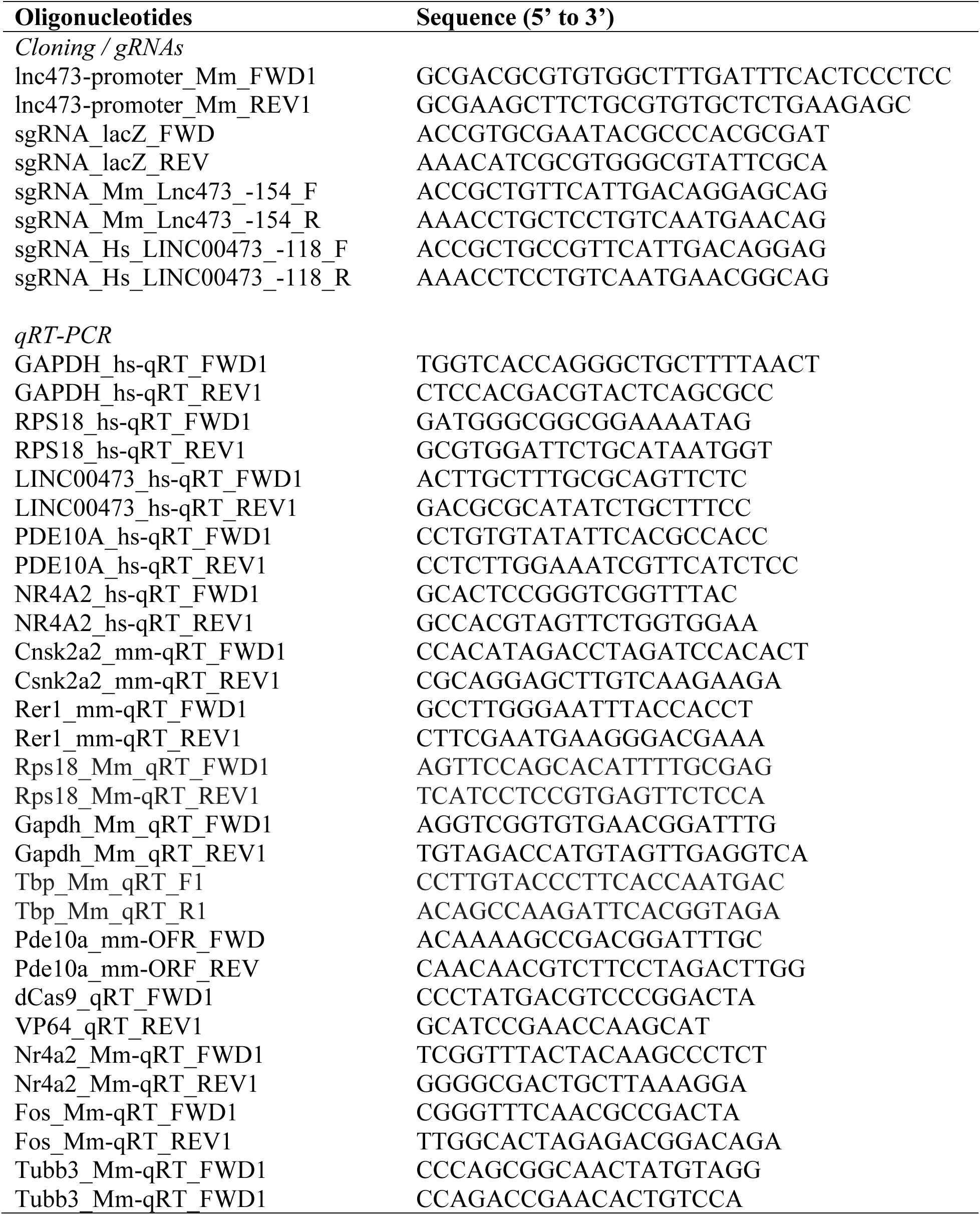
Oligonucleotide primer sequences used within this study.

